# Smaug regulates germ plasm synthesis and primordial germ cell number in Drosophila embryos by repressing the *oskar* and *bruno 1* mRNAs

**DOI:** 10.1101/2023.02.27.530189

**Authors:** Najeeb U. Siddiqui, Angelo Karaiskakis, Aaron L. Goldman, Whitby V.I. Eagle, Craig A. Smibert, Elizabeth R. Gavis, Howard D. Lipshitz

## Abstract

During Drosophila oogenesis, the Oskar (OSK) RNA-binding protein (RBP) determines the amount of germ plasm that assembles at the posterior pole of the oocyte. Here we identify the mechanisms that regulate the *osk* mRNA in the early embryo. We show that the Smaug (SMG) RBP is transported into the germ plasm of the early embryo where it accumulates in the germ granules. SMG binds to and represses translation of the *osk* mRNA itself as well as the *bruno 1* (*bru1*) mRNA, which encodes an RBP that we show promotes germ plasm production. Loss of SMG or mutation of SMG’s binding sites in the *osk* or *bru1* mRNAs results in ectopic translation of these transcripts in the germ plasm and excess PGCs. SMG therefore triggers a post-transcriptional regulatory pathway that attenuates germ plasm synthesis in embryos, thus modulating the number of PGCs.

## Introduction

In *Drosophila, C. elegans, Xenopus* and many other animals, a specialized region of the egg cytoplasm known as the germ plasm directs the formation and fate of the primordial germ cells (PGCs) of the early embryo (*1, 2*). Within the germ plasm reside electron-dense, membraneless organelles known as germ granules, which contain specific molecular determinants essential for PGC fate (*3*).

A genetic pathway has been defined in for the assembly of Drosophila germ plasm and germ granules (*1, 4, 5*). During oogenesis, *oskar* (*osk*) mRNA is transported to the posterior pole of the oocyte where it is translated. OSK protein nucleates assembly of the germ plasm by recruiting other germ plasm components, including Vasa (VAS) and Tudor (TUD).

Translation of *osk* mRNA is kept repressed during its transport to the posterior pole of the oocyte (*6*). Furthermore, localization of *osk* transcripts to the posterior is inefficient, with over 80% of the *osk* mRNA escaping localization (*7*), necessitating mechanisms to prevent this pool of mRNA from producing OSK protein. A major repressor of *osk* mRNA translation outside of the germ plasm is the Bruno 1 (BRU1) RNA-binding protein (*6, 8-10*), which binds to *cis*-elements (BRU1 response elements or BREs) in the *osk* 3’-untranslated region (3’UTR). BRU1 recruits the eIF4E-binding protein, Cup, which prevents translation initiation (*11, 12*). BRU1 also promotes the formation of complexes that silence translation of *osk* mRNA by preventing ribosome association (*10*) and, recently, has been shown to form a ‘scaffold’ that participates in a liquid-to-solid phase transition of *osk* mRNA granules (*13*). Unexpectedly, one of the clusters of BREs (the ‘C’ cluster) is also required for translation of the *osk* mRNA in the germ plasm; however, the early oogenesis arrest of *bru1* mutants has precluded proof that BRU1 is the *trans*-acting factor that acts through this cluster in the germ plasm (*9*).

The PGCs are the first cells to form in the early Drosophila embryo. They bud from the posterior pole when the nuclei arrive there after the eighth nuclear cleavage and undergo one or two rounds of division to give 30 to 35 PGCs, after which they migrate to the somatic component of the gonad (*1, 4, 5*). The number of PGCs that form depends on the quantity of germ plasm. Loss of germ plasm, as occurs in mutants for *osk* or other germ plasm components, results in failure to produce PGCs (*14-16*) while reduction of *osk* gene dosage from two copies to one results in about a 50% decrease in PGC number (*17*). In contrast, over-expression of OSK protein by increasing the gene dose to four or six copies results in a 25 to 50% increase in PGC number (*17, 18*).

In *Drosophila*, the maternally encoded RNA-binding protein, Smaug (SMG), is absent from ovaries but is synthesized in early embryos, where it directs translational repression and/or degradation of a large number of maternal mRNAs by binding to hairpin structures known as SMG recognition elements (*19-22*). Subsequently, SMG protein is cleared from the bulk cytoplasm (*23-25*) but not from the germ plasm and the PGCs (*26, 27*).

Here we show that SMG protein is transported into the germ plasm of early embryos where it accumulates and is integrated into the germ granules. Surprisingly, embryos from *smg*-mutant females have a 30% increase in the number of PGCs, which results from production of excess OSK in the germ plasm of the early embryo. We show that SMG represses translation of *osk* and *bru1* mRNAs in the germ plasm. Mutation of the two SREs in the *osk* mRNA or the five SREs in the *bru1* mRNA results in upregulation of OSK and BRU1 expression, respectively, and a greater number of PGCs than controls carrying functional SREs. Thus, SMG modulates PGC number by controlling the amount of both OSK and BRU1 in the germ plasm of early embryos. An implication of these results is that post-transcriptional regulation of germ plasm assembly at two distinct developmental stages determines PGC number: first, by germ plasm synthesis at the posterior pole of the oocyte and, second, by attenuation of germ plasm synthesis at the posterior pole of the embryo.

## Results

### SMG protein accumulates in the germ plasm of early embryos

Translation of *smg* mRNA is repressed in Drosophila oocytes but derepressed upon egg activation (*20*). SMG protein accumulates during the first two hours of embryogenesis and then is rapidly eliminated during interphase of nuclear cycle 14 *(23-25, 27, 28)*. An exception occurs in the germ plasm and PGCs, where SMG persists (*26, 27*). To visualize SMG protein accumulation in the germ plasm of live embryos, we produced transgenic flies that express either mCherry-SMG or Venus-SMG under control of the *smg* promoter, 5’ and 3’UTRs.

Live imaging showed that SMG protein became enriched in the germ plasm as early as nuclear cycle 5 (NC5, ∼40 minute after fertilization), prior to arrival of the syncytial nuclei at the posterior pole (Venus-SMG; Figure 1A; Movie S1). SMG-containing particles showed net movement towards the germ plasm, with a greater mean velocity towards (0.47 µm/second, n = 28) than away from (0.26 µm/second, n = 19) the posterior pole. When the nuclei arrived at the posterior pole (NC8), SMG became associated with the centrosomes, was taken up into the PGCs when they formed (NC9), and segregated into both daughter cells during PGC division (mCherry-SMG; Figure 1B; Movie S2).

**Figure 1.**
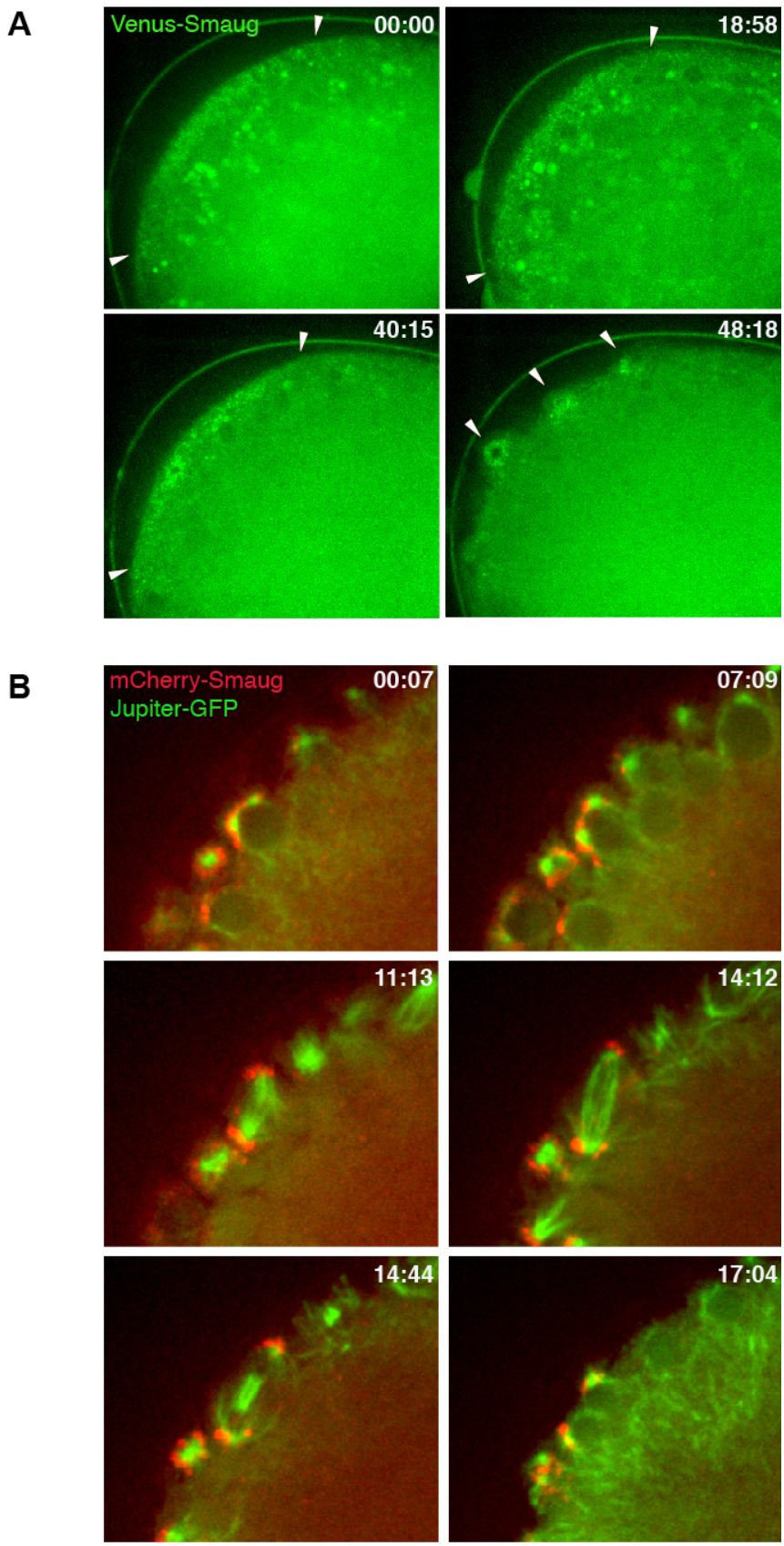
SMG protein is enriched in the germ plasm of early embryos. **(A)** Live imaging of Venus-SMG shows that SMG protein is enriched in the germ plasm as early as nuclear cycle (NC) 5. Still images were taken from Movie S1. The time after imaging began is indicated at the top right of each still image in minutes and seconds. White arrowheads indicate the boundaries of Venus-SMG in the germ plasm in the first three images; in the fourth image the arrowheads point to Venus-SMG associated with nuclei in budding PGCs. The transgene insertion was on the second chromosome (see Materials and Methods): *P{w*^*+*^*[Venus-SMG]:6}*. **(B)** Live imaging of mCherry-SMG (red) showing association of SMG with the spindle poles (green; detected with Jupiter-GFP) upon arrival of nuclei at the posterior (NC8) and incorporation into the PGCs when they bud (NC9). Still images were taken from Movie S2. The mCherry-SMG transgene insertion was on the second chromosome (see Materials and Methods): *P{w*^*+*^*[mCherry-SMG]:7}*.

### SMG is a component of the germ granules in embryos

It was previously shown that germ granules are transported on the astral microtubules of arriving nuclei about an hour after fertilization (*29, 30*). To assess whether SMG is a component of the germ granules we simultaneously imaged mCherry-SMG and the germ granule marker, VAS-GFP (*31*), in live embryos. We found that these proteins co-localize prior to, during, and after budding of the PGCs (Figure 2A; Movie S3). Immuno-electron microscopy confirmed that SMG and VAS co-localize in germ granules (Figure 2B, left and center panels).

**Figure 2.**
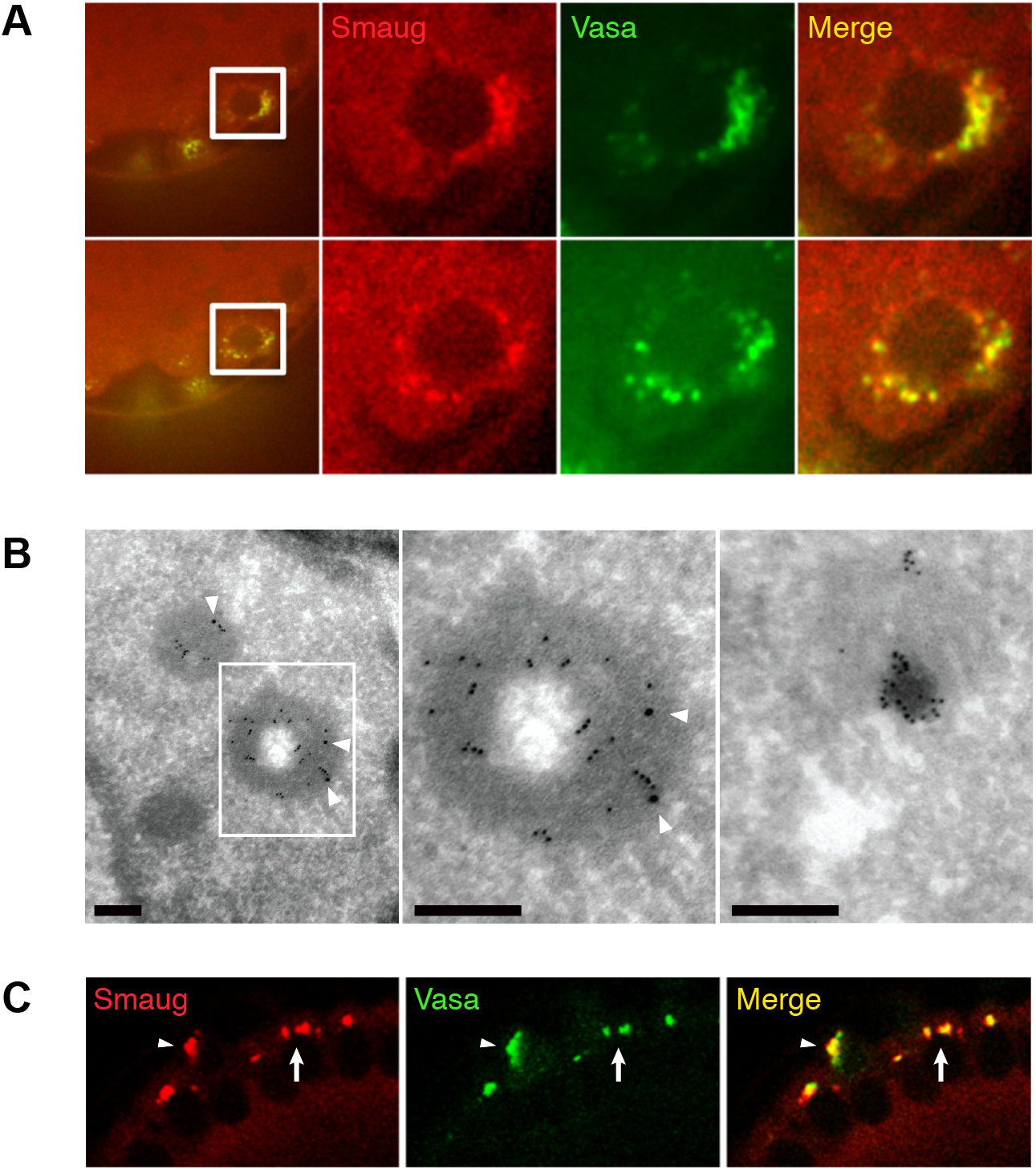
SMG is a component of the germ granules. **(A)** SMG co-localizes with VAS, a known component of germ granules. Panels are still images taken from movies of embryos co-expressing mCherry-SMG (red) and VAS-GFP (green). The boxed areas in the left-hand images are shown at higher magnification on the right. See associated Movie S3. The mCherry-SMG transgene insertion was on the third chromosome (see Materials and Methods): *P{w*^*+*^*[mCherry-SMG]:8}*. **(B)** Immuno-electron microscopy shows that, in the pole cells, SMG (10 nm gold: small particles) is found in germ granules along with VAS (15 nm gold: large particles, indicated with white arrowheads). The boxed area in the left-hand image is shown at higher magnification in the center. In the posterior somatic cells (right panel), SMG (10 nm gold) is enriched in electron-dense, non-membrane-bound organelles. Scale bars: 0.2 μm **(C)** SMG and VAS also co-localize in the apical region of the posterior soma, representing germ granules that are not taken up into the PGCs when they bud. White arrowheads point to co-localization in a PGC; white arrows point to co-localization in the apical somatic cytoplasm. See also associated Figure S1. The mCherry-SMG transgene insertion was on the second chromosome (see Materials and Methods): *P{w*^*+*^*[mCherry-SMG]:7}*.

A subset of SMG remained in the posterior cytoplasm, apical to the somatic nuclei well after budding of the PGCs was complete (Figure 2C). Its electron-dense, non-membrane-bound nature (Figure 2B, right panel) as well as co-localization of SMG protein and VAS protein (Figure 2C) indicated that this population of SMG represents germ plasm that is left behind after formation of the PGCs (*32*).

To assess whether VAS and SMG are dynamic components of the germ granules, we carried out fluorescence recovery after photobleaching (FRAP) analysis using VAS-GFP and mCherry-SMG (Figure S1). VAS-GFP rapidly recovered in the germ granules in the PGCs (Figure S1A) as well as in the granules that remain apical to the somatic nuclei (Figure S1B): The time to half-recovery for VAS in the PGC-located granules was ∼10 seconds and in the granules in the apical somatic cytoplasm was <5 seconds. This recovery rate is faster than that reported recently for VAS (33 seconds) (*33*) and the fractional recovery is somewhat higher. mCherry-SMG recovered slowly (75 seconds and 45 seconds for PGC granules and left-behind granules, respectively), and the fractional recovery was low (Figure S1C, D). We conclude that SMG is a component of germ granules with low mobility relative to VAS.

### *smg* mutants produce extra PGCs

Given the presence of SMG in the germ plasm of early embryos we next assessed whether removal of SMG had any consequences for the production of PGCs. We examined the original *smg* allele (*smg*^*1*^), which introduces a stop codon before the RNA-binding domain, resulting in synthesis of a truncated protein (*23, 28*), as well as a protein-null allele *(smg*^*47*^) (*19*). Two wild-type and four *smg*-mutant genotypes were assayed. Significantly more PGCs were found in embryos from *smg*-mutant females (henceforth referred to as ‘*smg*-mutant embryos’) than in wild type (Figure 3A): overall, the mean number of PGCs in the *smg* mutants was 40.2 versus 30.5 for wild type, a 32% increase.

**Figure 3.**
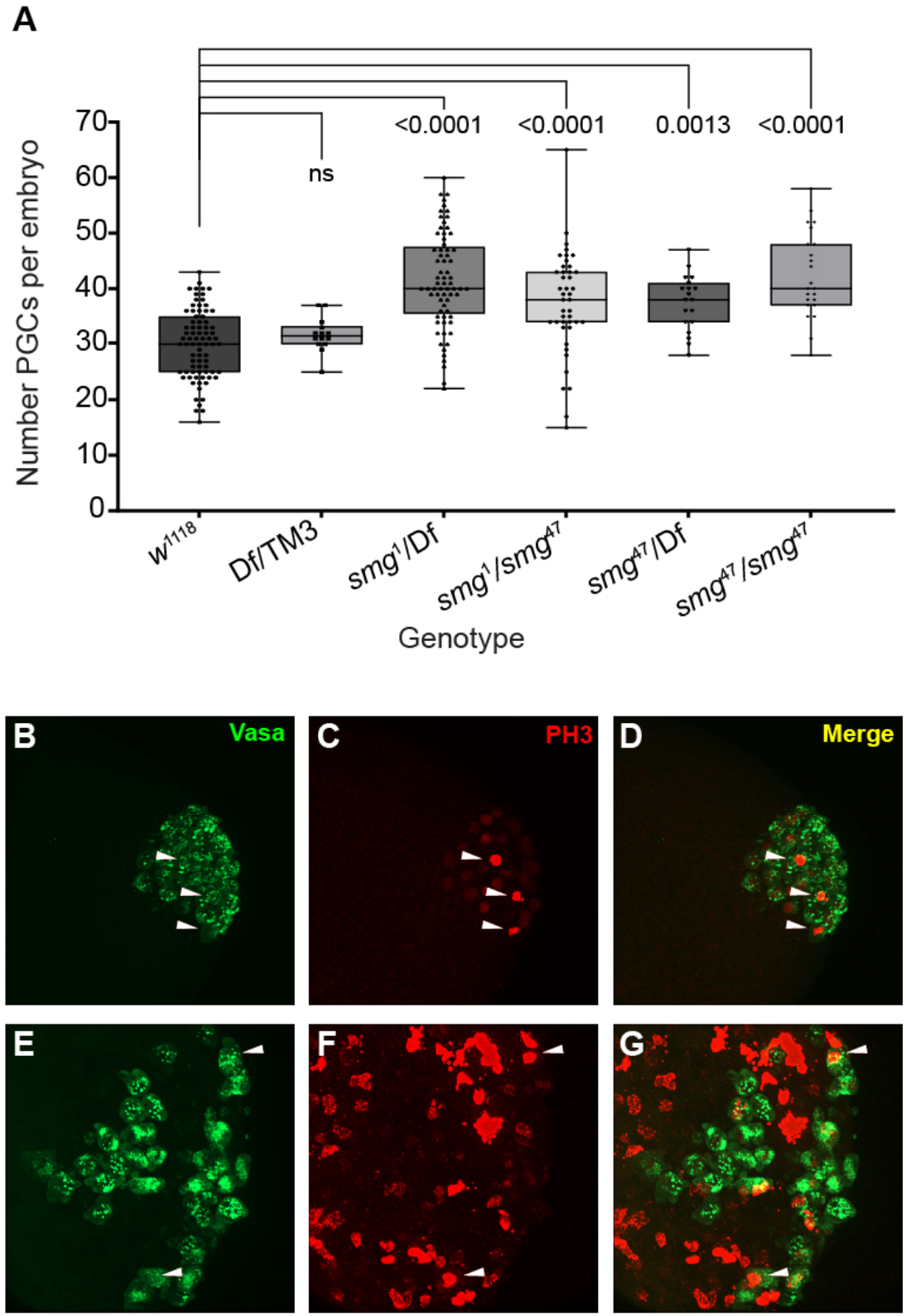
Extra PGCs form in *smg* mutants. **(A)** Box plots showing the number of PGCs in embryos from a variety of wild-type or *smg*-mutant mothers. *P* values are from the Kruskal Wallis One-Way ANOVA followed by Dunn’s test for multiple comparisons. **(B-G)** Confocal images showing VAS (green) and Phospho-histone H3 (PH3; red) in wild type (*w*^*1118*^) **(B, C, D)** or in *smg* mutants **(E, F, G)**. The *smg* mutant genotype of the females from which the embryos shown in **(E-G)** were obtained was *smg*^*1*^/*Df(3L)Scf-R6*. White arrowheads point to VAS-positive cells that are also PH3 positive. Note that in *smg* mutants the somatic nuclei continue to divide at this stage (*23*), and therefore are PH3-positive but VAS-negative. Three-dimensional reconstructions were used to distinguish the VAS-positive, PH3-positive PGC cells from the VAS-negative, PH3-positive somatic nuclei (see Materials and Methods).

There are two possible reasons – not mutually exclusive – for the appearance of extra PGCs in *smg* mutants: a greater number of PGCs may bud from the posterior tip of the embryo or the normal number may bud but then undergo more cell divisions than in wild type. To assess these possibilities, we performed immunofluorescence to visualize VAS, as a marker of PGCs, together with phosphohistone H3 (PH3), which marks cells in mitosis (*34*), in wild-type and *smg*-mutant embryos. The mean number of PH3-positive PGCs in wild type was 2.5 whereas it was 1.0 and 0.7 in *smg1/Df* and *smg*^*1*^*/smg*^*47*^, respectively (Figure 3B-G and Table S1; Bonferroni corrected Wilcoxon two-sided rank sum *P* = 0.003 and *P* = 10^−5^, respectively). We note that this is an underestimate of the difference since we did not normalize to PGC number, which is significantly higher in *smg* mutants than wild type (Figure 3A).

Since there are fewer – rather than more – dividing PGCs in *smg* mutants than in wild type, we conclude that more PGCs bud from the posterior pole of *smg*-mutant than wild-type embryos.

### SMG attenuates synthesis of OSK protein in the germ plasm of embryos

The increase in PGC number in *smg* mutants is similar to that previously observed in embryos from females carrying extra copies of the *osk* gene (*17, 18*). To assess whether, in *smg* mutants, there is a concomitant increase in *osk* mRNA and/or OSK protein levels in the germ plasm prior to budding of the PGCs, we performed single-molecule fluorescent *in situ* hybridization (smFISH) to detect *osk* mRNA as well as anti-OSK immunofluorescence. We found that more OSK is present in the germ plasm of *smg*-mutant embryos than in wild type prior to, as well as during, budding of the PGCs (Figure 4). Degradation of *osk* mRNA in the germ plasm proceeded normally in *smg* mutants; thus, there is a similar amount of *osk* mRNA in the germ plasm of wild-type and *smg*-mutant embryos (Figure S2). Since SMG protein is absent from oocytes and is not synthesized until after egg activation (*20, 23*), these data suggest that SMG attenuates translation of the *osk* mRNA in the germ plasm of the early embryo.

**Figure 4.**
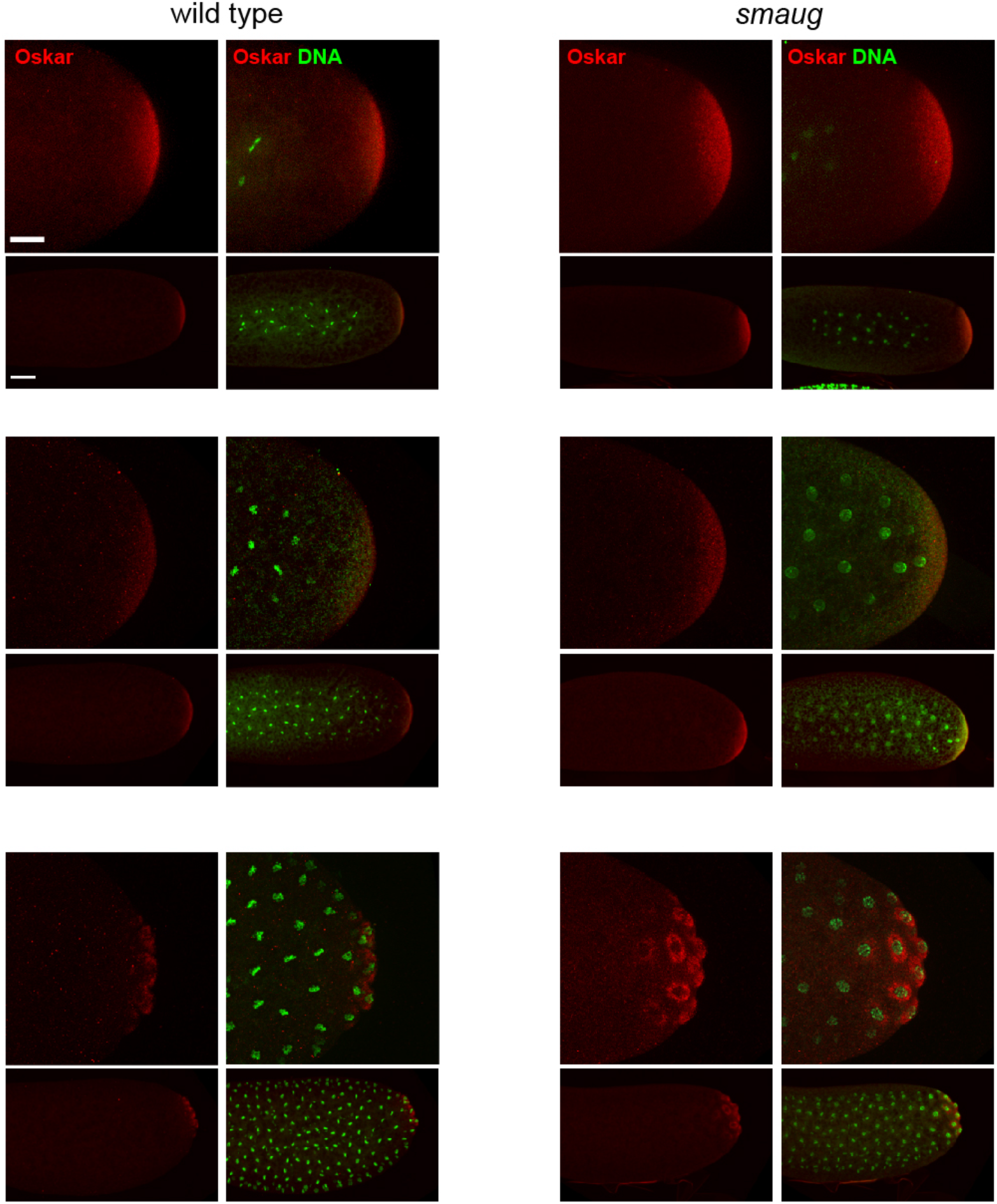
Excess OSK protein is produced in the germ plasm of *smg* mutants. Anti-OSK immunofluorescence in wild-type (left) or *smg*-mutant (right) embryos. The latter were from *smg*-mutant mothers (genotype: *smg*^*1*^/*Df(3L)Scf-R6)*. The top and middle sets of panels show the posterior pole of embryos prior to budding of the PGCs; the bottom set of panels shows the posterior pole of embryos during budding of the PGCs. Images are z-stack-projected confocal images of OSK (red) and DNA as visualized with pico-green (green). Scale bar in the low magnification image is 50 μm; that in the high magnification image is 25 μm. See also associated Figure S2.

### Mutation of SREs in the *osk* mRNA upregulates OSK levels and PGC numbers

SMG regulates its target mRNAs by binding to *cis*-elements known as SMG recognition elements (SREs) (*21, 22*). The *osk* mRNA contains two predicted SREs, one in the coding region and one in the 3’UTR (Figure S3A). If SMG directly regulates *osk* mRNA to attenuate production of OSK protein, then mutation of these SREs should result in increased levels of OSK relative to controls with wild-type SREs. We, therefore, mutated the SREs without altering the encoded amino-acid sequence in the context of an *osk* genomic transgene (*18*) (Figure S3A). We then compared OSK expression and PGC numbers in embryos from females carrying a transgene either with wild-type SREs, *osk*^2xSRE(+)^, or with both SREs mutated, *osk*^2xSRE(–)^, in an *osk* RNA-null background (Figure 5). Transgenic OSK protein encoded by both transgenes was localized to the germ plasm (Figure 5A).

**Figure 5.**
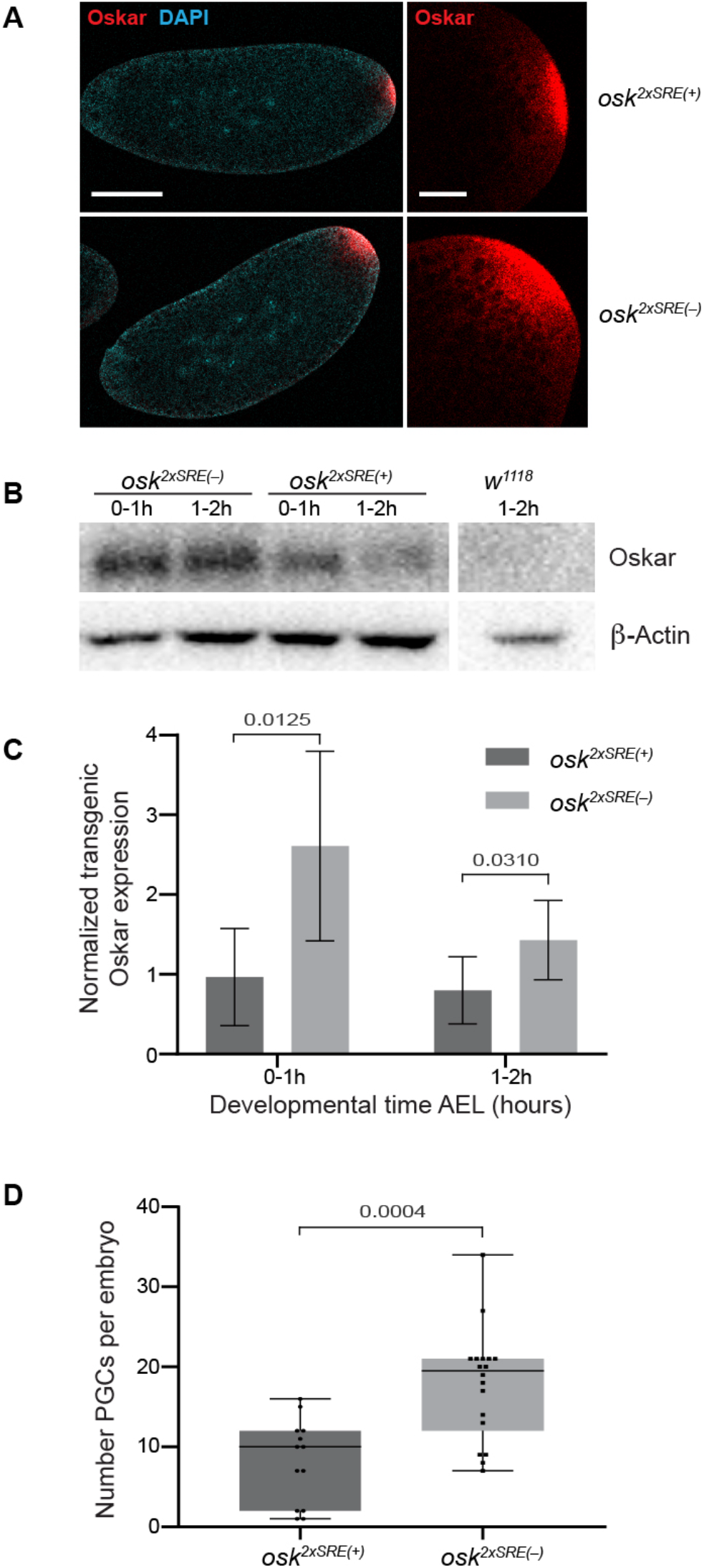
Mutation of SREs in the *osk* mRNA results in increased OSK protein and PGCs. **(A)** Confocal microscope images of NC 5 embryos expressing *osk*^2xSRE(+)^ or *osk*^2xSRE(–)^ in an *osk*^*0*^*/Df(osk)* background. Red: OSK, blue: DAPI. In both genotypes, OSK is localized to the germ plasm. Scale bar in the low magnification image is 100 μm; that in the high magnification image is 25 μm. **(B)** A representative western blot of embryos expressing *osk*^2xSRE(+)^ or *osk*^2xSRE(–)^ in an *osk*^*0*^*/Df(osk)* background. **(C)** Quantification of OSK in the two genotypes described in (B). Values were normalized to the actin loading control and then to *osk*^2xSRE(+)^ at the 0-1 hr time point. n = 5. **(D)** Box plots of PGC numbers in embryos expressing *osk*^2xSRE(+)^ or *osk*^2xSRE(–)^ in an *osk*^*0*^*/Df(osk)* background. Each dot or square represents the PGC count in a single embryo. *P* values in (C) and (D) are from the one-sided Wilcoxon rank sum test.

The amount of OSK in both 0-1 and 1-2 hr embryos from *osk*^2xSRE(–)^ females was significantly higher than in embryos from *osk*^2xSRE(+)^ females (Figure 5B, C; Wilcoxon one-sided rank sum test *P* = 0.0125 and 0.031, respectively, n = 5). Both transgenes gave only partial rescue of PGC number, possibly because the C-terminal epitope tags somewhat compromise OSK function. This sensitized the rescue experiment, leading to strikingly more PGCs in *osk*^2xSRE(–)^ embryos (approximately double the number) than in *osk*^2xSRE(+)^ embryos (Figure 5D; n = 13 and 18, respectively; Wilcoxon one-sided rank sum test *P* = 0.0004). Together these data are consistent with direct regulation of *osk* mRNA by the SMG RNA-binding protein to repress *osk* translation in the germ plasm of embryos, thus attenuating PGC numbers.

### SMG represses translation of *bru1* mRNA in the germ plasm

The BRU1 RNA-binding protein plays a key role in assembly of *osk* RNA-containing granules during oogenesis (*10, 13*). BRU1 was initially identified as a translational repressor of *osk* transcripts in the bulk cytoplasm of the oocyte (*6, 8*). Subsequent experiments suggested that, in contrast to the bulk cytoplasm, BRU1 might function in positive regulation of *osk* mRNA translation in the oocyte germ plasm (*9*).

In embryos, the *bru1* mRNA is cleared from the bulk cytoplasm but *bru1* transcripts are enriched in the germ plasm and are taken up into the PGCs when they bud (*26, 35*). In our genome-wide study, where we purified SMG from early embryos together with its bound RNAs, *bru1* mRNA was a top hit (*19*).

To assess the spatial aspects of SMG’s association with *bru1* mRNA in the germ plasm, we carried out single-molecule fluorescent *in situ* hybridization (smFISH) of *bru1* mRNA in Venus-SMG-expressing embryos; as controls, we also monitored two of SMG’s previously identified target mRNAs, *Hsp83* and *nanos* (*nos*) (Figure S4). As SMG became enriched in the germ plasm, there was significant co-localization of SMG foci and *bru1* transcripts.

To assess whether SMG might repress translation of the *bru1* mRNA in the germ plasm we analyzed BRU1 protein in wild-type and in *smg*-mutant embryos by immunofluorescence. In wild-type embryos, BRU1 was very weakly detectable in the early embryonic germ plasm and in the PGCs (Figure 6). In contrast, in *smg*-mutant embryos, BRU1 was readily detectable and increased over time in the germ plasm (Figure 6). These data are consistent with the hypothesis that SMG represses the translation of *bru1* transcripts in germ plasm.

**Figure 6.**
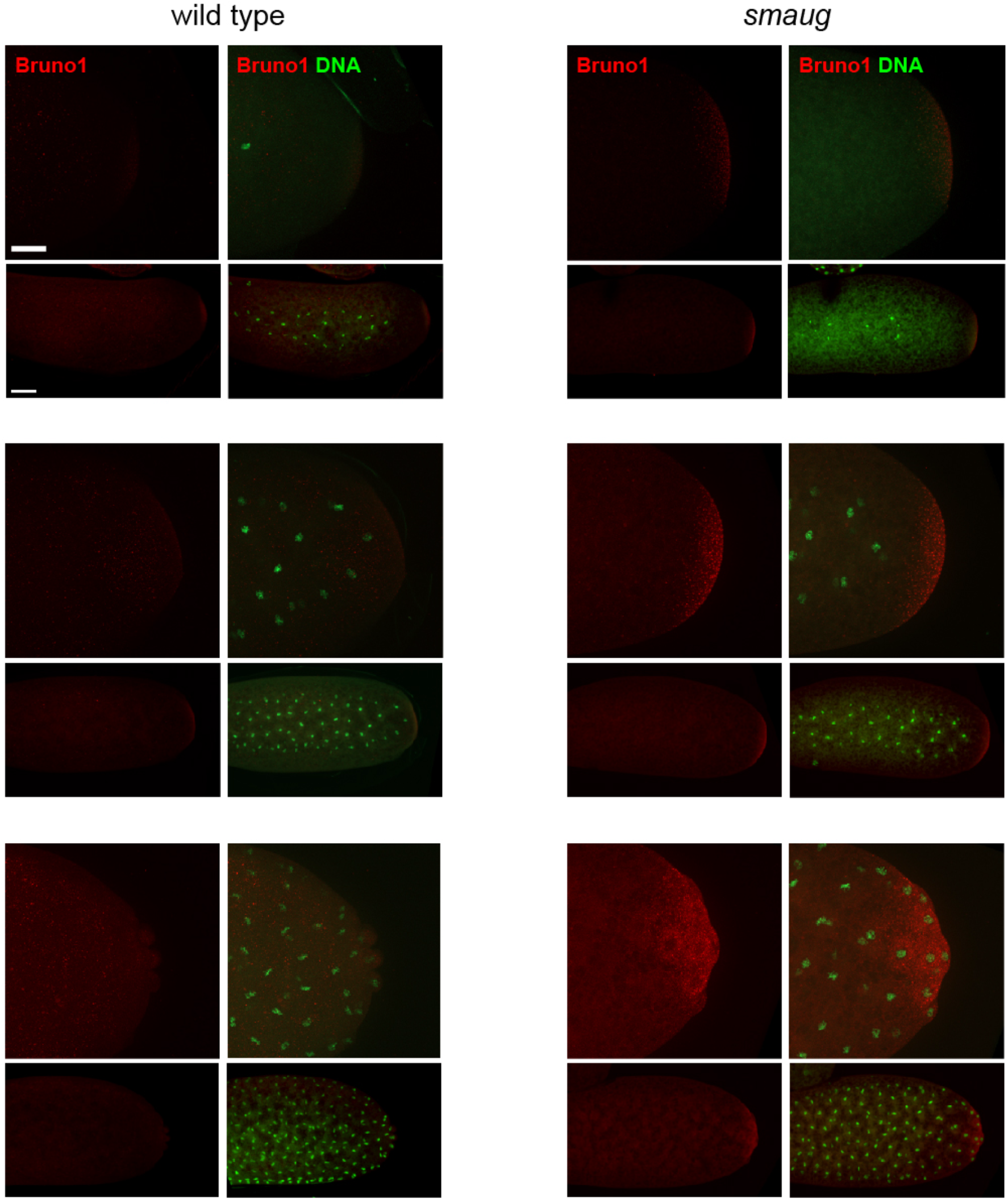
SMG represses *bru1* mRNA in the germ plasm. Anti-BRU1 immunofluorescence in wild-type (left) or *smg*-mutant (right) embryos. The latter were from *smg*-mutant mothers (genotype: *smg*^*1*^/*Df(3L)Scf-R6*). The top and middle sets of panels show the posterior pole of embryos prior to budding of the PGCs and the bottom set of panels shows the posterior pole of embryos during budding of the PGCs. Images are z-stack-projected confocal images of BRU1 (red); DNA as visualized with pico-green (green). Scale bar in the low magnification image is 50 µm; that in the high magnification image is 25 µm.

### Mutation of SREs in the *bru1* mRNA upregulates germ plasm BRU1 levels and PGC numbers

The *bru1* mRNA contains five predicted SREs, all in its coding region (Figure S3B). If SMG binds to and translationally represses *bru1* mRNA, then mutation of these SREs should result in synthesis of excess BRU1 in the embryo’s germ plasm. To test this, we generated transgenes in which the *bru1* open reading frame (ORF) and 3’UTR were placed under the control of the UASp promoter (*36*). For one, we used a wild-type *bru1* ORF while, for the other, we mutated the five predicted SREs without altering the amino acid sequence of the encoded BRU1 protein (Figure S3B). The mRNAs encoded by these transgenes, abbreviated *bru1*^*5xSRE(+)*^ and *bru1*^*5xSRE(–)*^, were expressed during oogenesis driven by maternal tubulin-GAL4 (*37, 38*).

Confocal immuofluorescence analysis showed that the BRU1 protein in both *bru1*^*5xSRE(–)*^ and *bru1*^*5xSRE(+)*^ embryos was enriched in the germ plasm (Figure 7A). Western blots showed that the levels of BRU1 in 0-1 and 1-2 hour embryos were higher in *bru1*^*5xSRE(–)*^ than *bru1*^*5xSRE(+)*^ embryos (Figure 7B,C; Wilcoxon one-sided rank sum test *P* = 0.0896 and 0.002, respectively, n = 4). These data are consistent with the hypothesis that SMG directly represses translation of the *bru1* mRNA in the germ plasm of early embryos.

**Figure 7.**
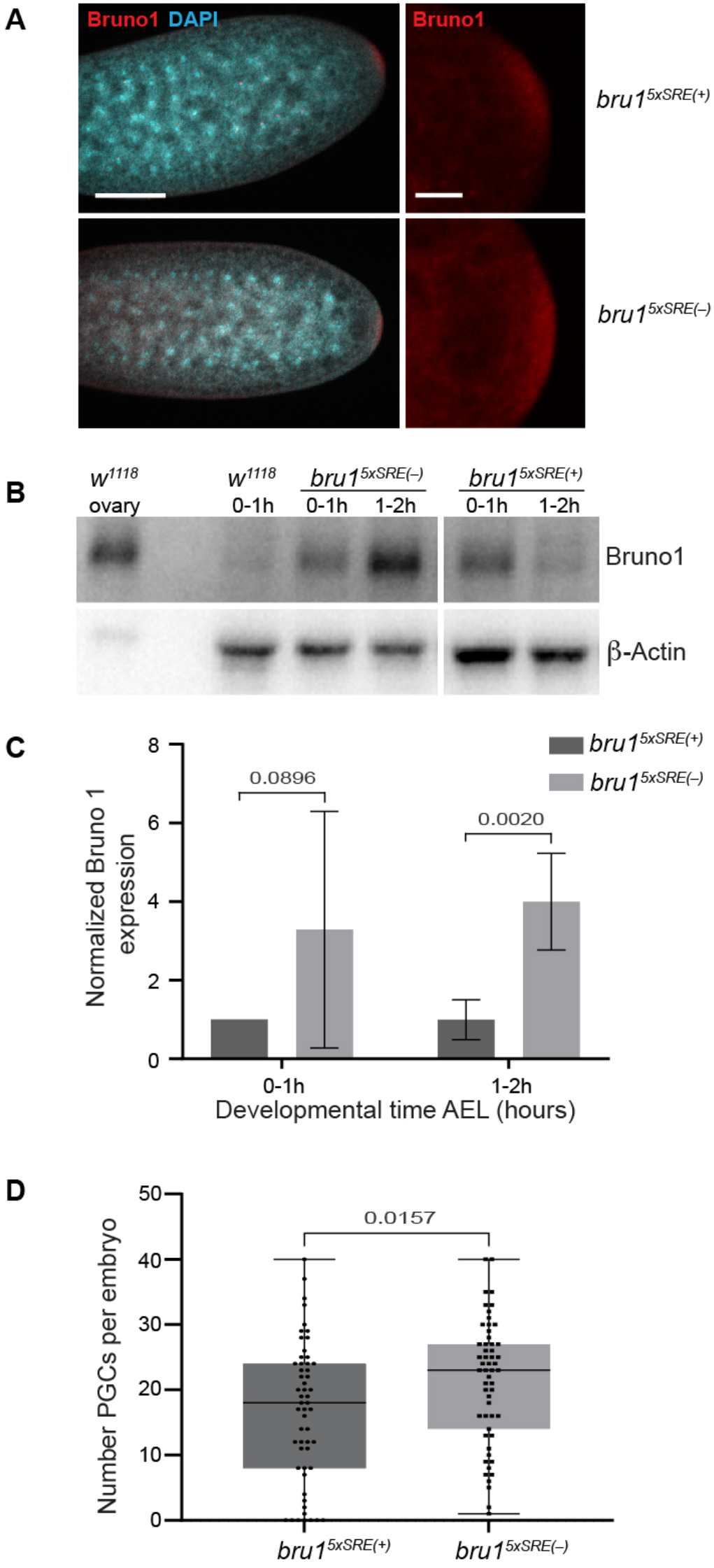
Mutation of SREs in the *bru1* mRNA results in increased BRU1 protein and PGCs. **(A)** Confocal microscope images of NC 7 embryos expressing *bru1*^5xSRE(+)^ or *bru1*^5xSRE(–)^. Red: BRU1, blue: DAPI. In both genotypes, BRU1 is localized to the germ plasm. Scale bar in the low magnification image is 100 µm; that in the high magnification image is 25 µm. **(B)** A representative western blot of embryos expressing *bru1*^5xSRE(+)^ or *bru1*^5xSRE(–)^ in a wild-type background. **(C)** Quantification of BRU1 in the two genotypes described in (B). Values were normalized to the actin loading control and then to *bru1*^5xSRE(+)^ at the 0-1 hr time point. n = 4. **(D)** Box plots of PGC numbers in embryos expressing *bru1*^5xSRE(+)^ or *bru1*^5xSRE(–)^. Each dot or square represents the PGC count in a single embryo. *P* values in (C) and (D) are from the one-sided Wilcoxon rank sum test.

We next asked whether the increase in BRU1 levels in embryos expressing *bru1*^*5xSRE(–)*^ mRNA resulted in a greater number of PGCs than embryos expressing the *bru1*^*5xSRE(+)*^ mRNA. UAS-driven over-expression of BRU1 during oogenesis is known to reduce posterior OSK levels and the amount of germ plasm produced during oogenesis (*39*). Consistent with this earlier observation, NC 14 embryos from females that expressed transgenic *bru1* transcripts driven by the maternal tubulin-Gal4 driver had fewer pole cells than wild type. Strikingly, embryos from females expressing *bru1*^*5xSRE(–)*^ had significantly more PGCs than those expressing *bru1*^*5xSRE(+)*^ mRNA (Figure 7D; n = 55 each; Wilcoxon one-sided rank sum test *P* = 0.0157). This result is consistent with the BRU1 synthesized in the germ plasm of *bru1*^*5xSRE(–)*^ embryos rescuing the reduced number of pole cells caused by over-expression of BRU1 during oogenesis to a significantly greater extent than the lower BRU1 levels in *bru1*^*5xSRE(+)*^ embryos.

Together these results are consistent with the hypothesis that BRU1 can potentiate production of germ plasm and PGCs in early embryos, and that SMG-directed repression of both OSK and BRU1 synthesis in early embryos attenuates germ plasm production.

## Discussion

While there have been numerous studies of germ plasm synthesis during oogenesis, ours is the first to focus on post-fertilization regulatory mechanisms that modulate the amount of germ plasm in the embryo. We have shown that the SMG RNA-binding protein, which is synthesized only after fertilization, sits at the top of a post-transcriptional cascade that attenuates germ plasm synthesis in the embryo and, consequently, modulates the number of PGCs (Figure 8). Notably, SMG regulates two key germ plasm components: First, SMG directly binds and represses *osk* transcripts; second, SMG binds and represses *bru1* transcripts, which encode a positive regulator of *osk* translation in the germ plasm. Abrogation of SMG binding to either of these target RNAs results in upregulation of germ plasm and PGC numbers.

**Figure 8.**
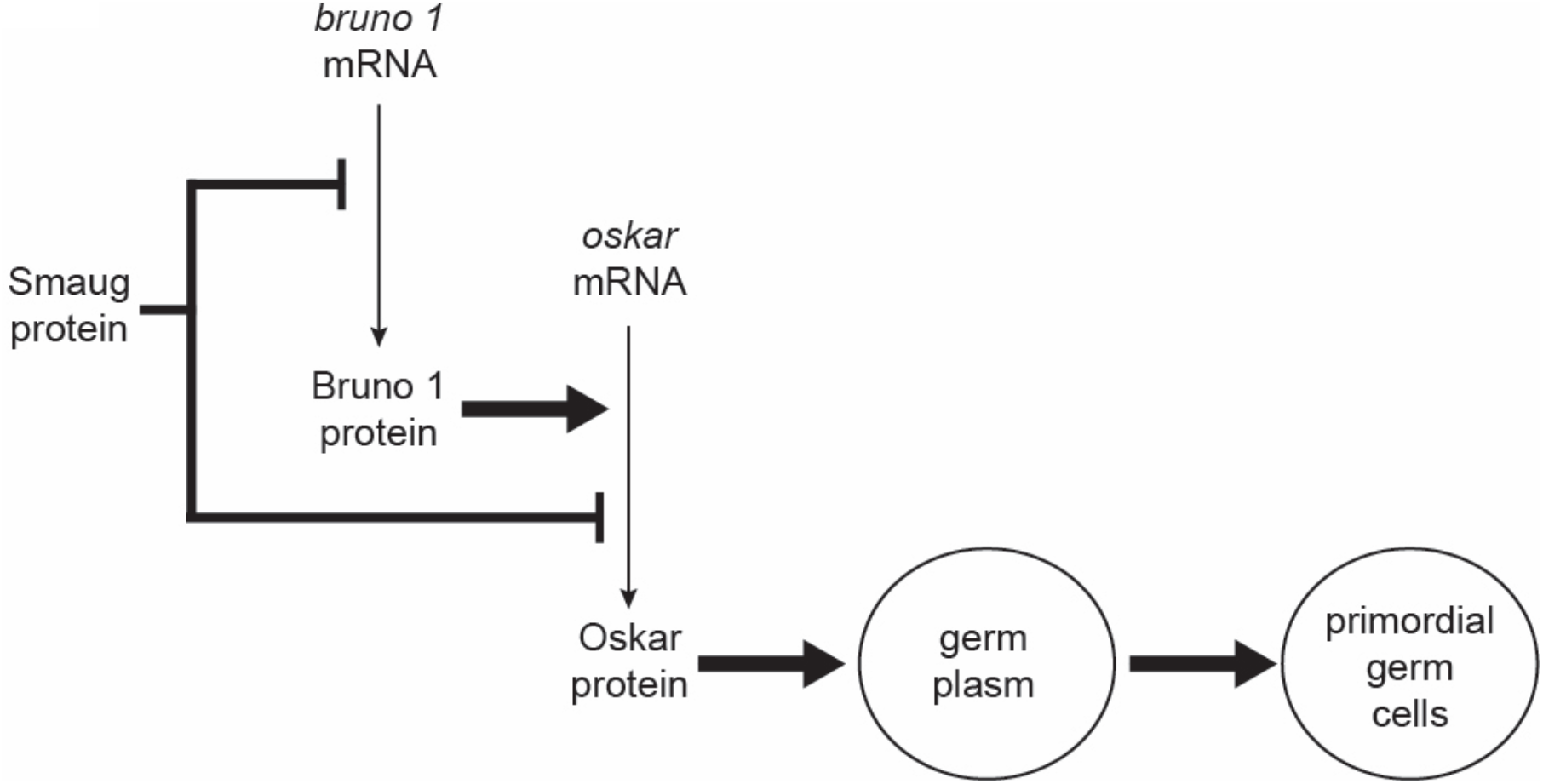
Pathway for regulation of PGC number in embryos. SMG protein is synthesized and transported into the germ plasm of the early embryo. There SMG represses translation of the *osk* and *bru1* mRNAs thus attenuating synthesis of OSK protein and modulating PGC number.

The exact location within the germ plasm where SMG implements repression of the *osk* and *bru1* mRNAs remains to be determined. The *osk* mRNA is not an integral component of mature germ granules but is present in ‘founder’ granules in the germ plasm within which *osk* transcripts undergo degradation (*30, 40*). In contrast, since both SMG protein and *bru1* mRNA are components of mature germ granules, this is the most likely site of SMG’s repression of *bru1*. Notably, of the three clusters of BREs in the *osk* 3’UTR, Cluster C was previously shown to participate in translational activation of *osk* mRNA in the germ plasm of late-stage oocytes (*9*). However, since *bru1* mutants arrest oocyte development early in oogenesis – indeed, the *bru1* gene was initially named *arrest* because of this (*41*) – it was not possible in the 2010 study to determine whether the activation of *osk* is directed by BRU1 itself or by a different RNA-binding protein that recognizes BREs. Our experiments have shown that BRU1 is indeed a positive regulator of *osk* mRNA translation and that it is capable of potentiating OSK synthesis in the embryo’s germ plasm.

Smaug-dependent repression of target transcripts in the bulk cytoplasm of the early embryo is accomplished via the eIF4E-binding protein, Cup (*42*), and the Argonaute 1 (AGO1) protein (*43*). These are, therefore, candidates to mediate SMG’s repression of the *osk* and *bru1* mRNAs in the germ plasm. Cup has been shown to be enriched in the germ plasm in both oocytes and early embryos (*12, 44*) where it physically interacts and co-localizes with OSK (*12, 44, 45*). Maternal reduction of Cup gene dose by 50% results in reduced PGC numbers in embryos, likely because of defects in germ plasm assembly and maintenance during oogenesis (*44*). There have not been any studies of the role of Cup in the embryonic germ plasm.

AGO1 also colocalizes with OSK at the posterior of oocytes (*46*) but AGO1 localization to the germ plasm of embryos has not been reported and we did not detect AGO1 in our earlier definition of the PGC proteome (*26*). Consistent with the possibility that AGO1 is, indeed, present and functional in the embryonic germ plasm and/or PGCs, reducing the dose of the *ago1* gene can suppress the mutant phenotype of the *polar granule component* (*pgc*) gene, which encodes a protein that is expressed transiently in PGCs after they bud and is essential for their survival (*47*).

Another Argonaute-family protein, Aubergine (AUB), is a known germ granule component (*48*) that directs degradation of *osk* mRNA in ‘founder’ granules (*40*) and was identified as a component of the PGC proteome (*26*). However, unlike AGO1, AUB has not been implicated in SMG’s repressive mechanism. Removal of AGO1, AUB or Cup specifically from the embryonic germ plasm will be required to assess whether extra PGCs form as in *smg* mutants.

Both *bru1* and *nos* mRNAs are known targets of SMG; however, *bru1* mRNA is repressed while *nos* mRNA is translated in the germ plasm. How SMG represses translation of *bru1* but not *nos* mRNA in the germ plasm remains to be elucidated. Two lines of evidence, not mutually exclusive, point towards possible mechanisms. First, smFISH together with super-resolution microscopy has shown that mRNAs are spatially organized into ‘homotypic clusters’ within the germ granules (*30, 33, 49, 50*). If SMG is associated with *bru1* clusters but not *nos* clusters then the latter transcripts would be translated but the former repressed. Second, OSK protein binds to *nos* mRNA (*51, 52*) and may function to abrogate SMG’s ability to repress *nos* translation in the germ plasm (*22, 28, 53, 54*). It has been shown that, in addition to its SREs, the *nos* mRNA is likely to contain one or more *cis*-elements that relieve translational repression in the germ plasm (*7, 21, 22*). We speculate that the *bru1* mRNA may lack these additional *cis*-elements and, therefore, translation of the *bru1* mRNA is repressed by SMG in the germ plasm.

The *osk* mRNA is robustly translated at the posterior pole of the oocyte. Since *osk* transcripts are present in the germ plasm of early embryos, in principle synthesis of OSK could continue after fertilization were SMG not present to repress translation of *osk* mRNA. What, then, is the biological significance of attenuation of germ plasm synthesis in embryos since, intuitively, one might think that production of more PGCs is advantageous? We speculate that the attenuation mechanism has evolved in *Drosophila* because of the intimate relationship between the germ plasm and anterior-posterior axis formation in the embryo. Body pattern defects have been shown to result when the dose of the *osk* gene and, thus, the amount of germ plasm, is increased (*17, 18*). This is a consequence of production of ectopic NOS protein in the anterior of embryos, causing repression of the *bicoid* mRNA, translation of which is essential for anterior pattern. Thus, the mechanism that we have uncovered in this study may have evolved, not specifically to attenuate the number of PGCs but, rather, to ensure correct body pattern. An implication is that, during the evolution of organisms such as *Drosophila* – where the germ plasm mechanistically links PGC formation and embryonic patterning – a balance has been achieved that optimizes the amount of germ plasm to ensure that both developmental processes can proceed unhindered. By attenuating germ plasm synthesis in the early embryo, SMG may achieve this balance.

## Materials and Methods

### Experimental Design

**Table 1:**
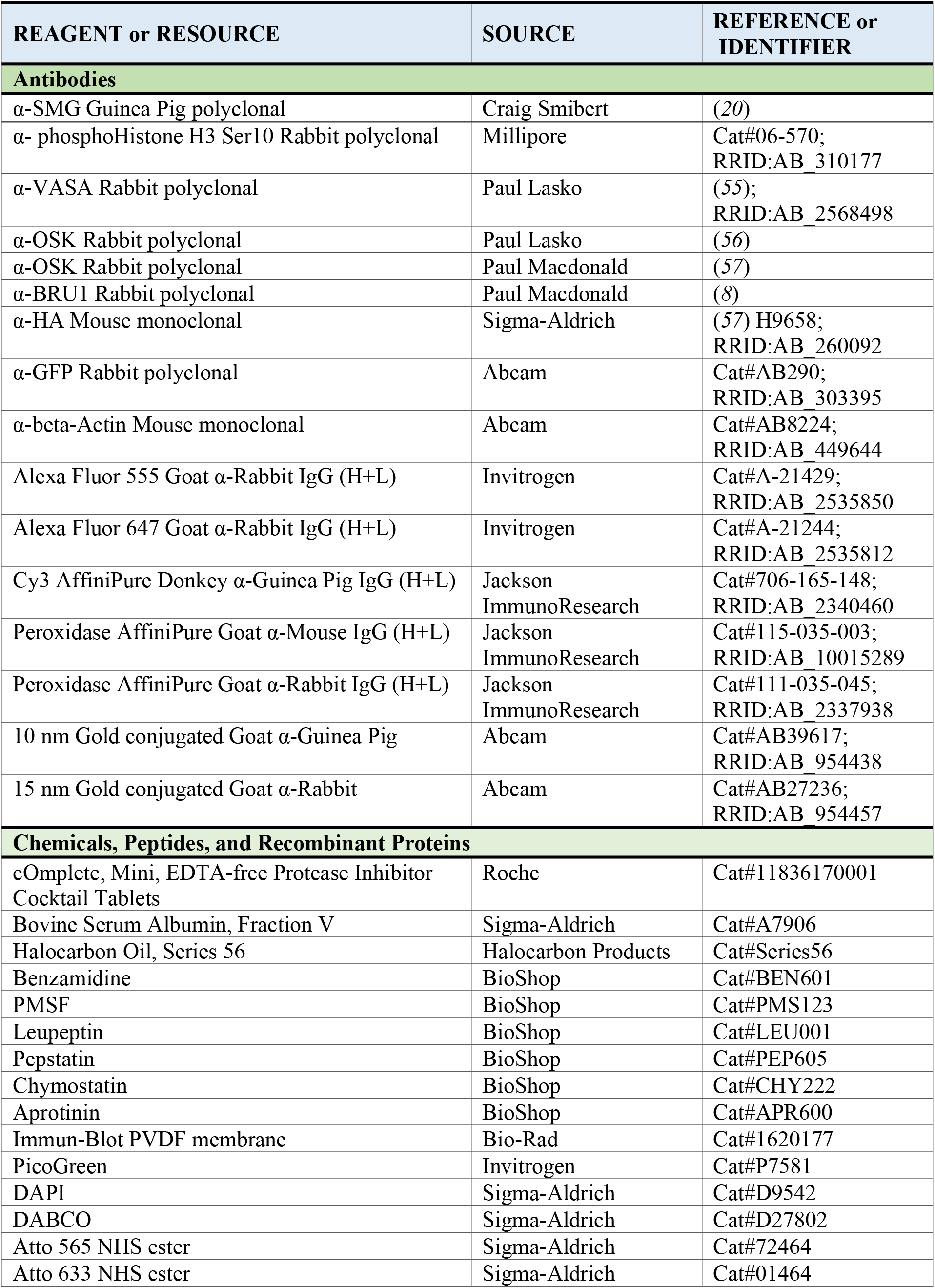
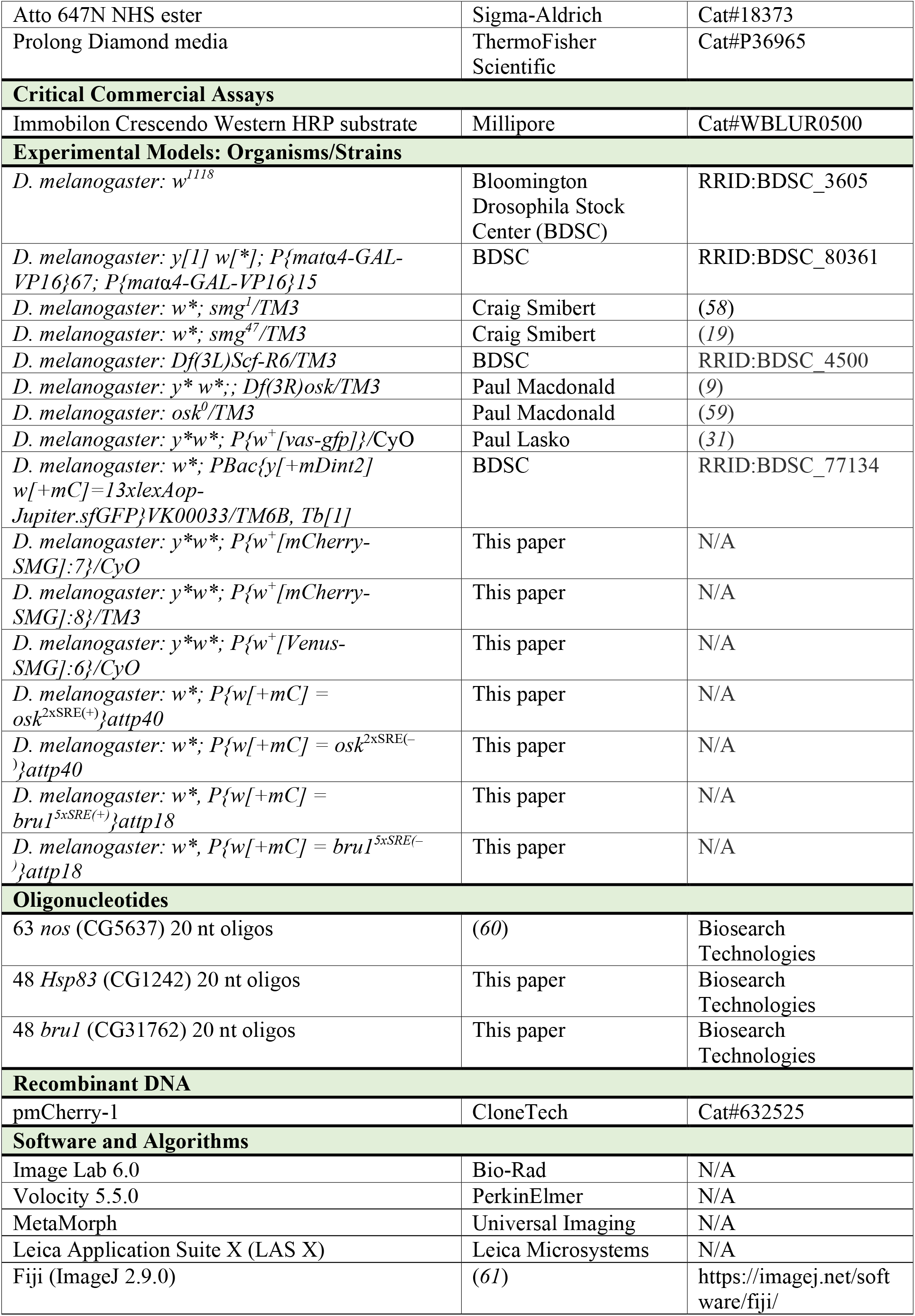
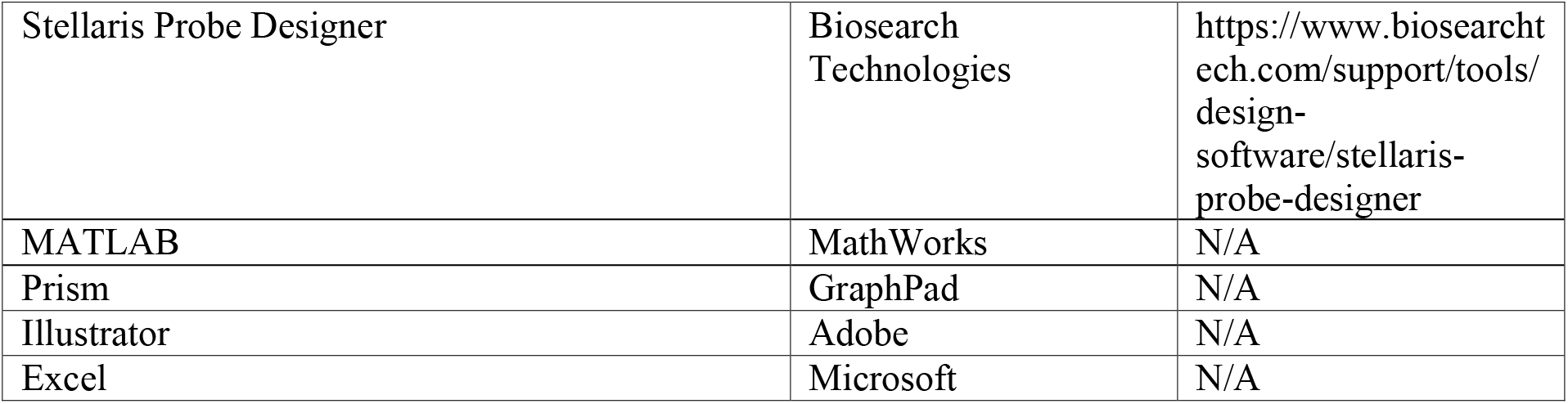
List of Reagents and Resources

### Drosophila culture and mutants

All stocks were maintained at 25°C, 65% humidity and a 12-hour day/12-hour night cycle on standard medium of cornmeal, yeast, agar, and molasses. The following fly strains were used: *w*^*1118*^ (Bloomington Drosophila Stock Center, BDSC #3605), *VAS-GFP* (*31*), *smg*^*1*^ (*28*), *smg*^*47*^ (*19*), *Df(3L)Scf-R6* (BDSC #4500), *Df(3R)osk* (*9*) and *osk*^0^ (*59*), *Jupiter-GFP* (BDSC #77134), *mCherry-SMG* (see below), *Venus-SMG* (see below). The *mCherry-SMG* and *Venus-SMG* transgenes completely rescue the *smg*^*1*^ mutant phenotype. *UASp* transgenes were expressed by crossing to the *P(matα4-GAL-VP16)67*; *P(matα4-GAL-VP16)15* driver (*37, 38*). Details on the generation of fly strains expressing the *osk*^2xSRE(+)^, *osk*^2xSRE(–)^, *bru1*^*5xSRE(+)*^ and *bru1*^*5xSRE(–)*^ transgenic constructs are given below.

### Construction of transgenes and production of transgenic flies

The N-terminal mCherry-SMG and Venus-SMG constructs were generated by replacing the N-terminal TAP-tag from the TAP-tagged SMG plasmid (*62*) with either mCherry or Venus tags based on the published sequences (*63, 64*). The resulting plasmid was digested with *Not*I and the *smg* genomic fragment with its regulatory regions and an N-terminal mCherry- or Venus-tag was subcloned into pCaSperR4 (*65*). Transgenic flies were generated by random - element mediated insertion of the transgene in the germ line.

The *osk*^2xSRE(+)^ and *osk*^2xSRE(–)^ transgenes contain an 8 kb genomic *osk* rescue fragment (*18*) cloned into *pCaSpeR*4-*attB* (see above). The original construct was modified to carry a *Kpn*I-*Bam*H1 fragment between the stop codon and the 3’UTR, which encodes 6xHis followed by 3xHA epitope tags (<u>ggt acc</u> ggc cgc gga tcg cat cac cat cac cat cac ggc cgc atc ttt tac cca tac gat gtt cct gac tat gcg ggc tat ccc tat gac gtc ccg gac tat gca gga tct tat cca tat gac gtt cca gat tac gct gct ta<u>g gat cc</u>; the underlined nucleotides are the *Kpn*I and *Bam*H1 sites, respectively). This transgene is referred to as *osk*^2xSRE(+)^. In *osk*^2xSRE(–)^ the predicted SRE sequences were altered by QuickChange mutagenesis (Figure S3A). Transgenic flies were generated by insertion of the transgene into the *phiC31 attP40* landing site on the second chromosome (*66*).

To produce the *UAS-bru1 5’UTR-bru1ORF*^*5xSRE+*^*-bru1 3’UTR* transgene (referred to as *bru1*^*5xSRE(+)*^) the *bru1* cDNA, P151 (a gift from Paul Macdonald), was cloned into a UASp vector (*36*) into which the *attB* site had been inserted. For the *UAS-bru1 5’UTR-bruORF* ^*5xSRE-*^*-bru1 3’UTR* construct (referred to as *bru1*^*5xSRE(–)*^), SRE mutations were introduced into the *bru1* ORF (using the *bru1* cDNA as a template) by PCR with overlapping primers that carried the mutations (Figure S3B). The mutated cDNA was cloned into the *UASp-attB* vector. Transgenic flies were generated by insertion of the transgene into the *phiC31 attP18* landing site on the X chromosome (*66*).

### Confocal immunofluorescence microscopy

Zero to four-hour (hr) old embryos were collected from cages on apple juice plates, dechorionated with 4.2% sodium hypochlorite for 2 minutes, and then rinsed with dH2O. Dechorionated embryos were fixed and permeabilized in 4% formaldehyde, 1xPBS and heptane for 20 min, and devitellinized by addition of methanol followed by vigorous shaking for 30 s. Fixed embryos were rehydrated by washing 4 times with PBSTx (1xPBS, 0.1% Triton X-100) and blocked with 10% bovine serum albumin (BSA) in PBSTx for 1 hr at room temperature. Embryos were incubated with rocking at 4°C overnight with primary antibodies diluted in 1% BSA in PBSTx. Embryos were washed 3 times for 15 min each with PBSTx at room temperature, then incubated with rocking in secondary antibodies diluted in 1% BSA in PBSTx for 1 hr at room temperature. Embryos were washed 5 times for 10 min each with PBSTx, and mounted in 2.5% DABCO, 70% glycerol in PBS.

Primary antibodies used for immunofluorescence were: guinea-pig anti-SMG (1:500) (*20*); rabbit anti-GFP (1:500) (Abcam, Inc.); rabbit anti-PhosphoHistone H3 Ser10 (1:200) (Upstate USA, Inc.); rabbit anti-VAS (1:10,000) (*55*); rabbit anti-OSK (1:250) for Figure 4 (*6*); rabbit anti-OSK (1:1,000) for Figure 5 (*56*); rabbit anti-BRU1 (1:500) (*8*). Secondary antibodies were: Cy3-conjugated donkey anti-guinea pig (1:300) (Jackson ImmunoResearch), goat anti-rabbit Alexa Fluor 555 (1:300) (Invitrogen), or goat anti-rabbit Alexa Fluor 647 (1:300) (Invitrogen). Picogreen (1:300) (Invitrogen) or DAPI (1:4000) (Sigma-Aldrich) were used to stain DNA.

Samples were visualized either with an Olympus IX81 inverted fluorescence microscope equipped with a Hamamatsu Back-Thinned EM-CCD camera (9100-13) and Quorum spinning disk confocal scan head or with a Leica SP8 inverted scanning confocal microscope. The acquired fluorescent images were imported into Volocity 5.5.0 software (PerkinElmer), or into Fiji (ImageJ 2.9.0, National Institutes of Health), respectively, and minimally processed, then cropped for figures in Illustrator (Adobe). When required, a single plane was obtained from approximately the middle of the embryo.

### Live imaging and analysis of SMG particle movement

For live imaging, embryos were collected for 15 min to 1 hr, depending on the stage to be imaged, dechorionated with 4.2% sodium hypochlorite for 2 minutes, washed with dH2O, mounted as a monolayer of embryos in a drop of halocarbon oil (Series 56) on the outer surface of the Greiner Lumox culture dishes (Sigma-Aldrich Cat# Z376744), and covered with a coverslip before imaging. All movies were captured using a 40x Planapochromat objective (NA 1.5) on a Nikon Eclipse TE2000-E inverted microscope with a Perkin-Elmer spinning disk and a Hamamatsu Orca-EM CCD camera. Movies were processed using the Metamorph software package (Universal Imaging, Downington, Pennsylvania).

### Cryo-immunogold electron microscopy

Embryos were collected 140-160 min after egg-laying, dechorionated with 4.2% sodium hypochlorite for 2 min, rinsed in dH2O, and then shaken for 30 min in a biphasic solution containing 8mL of heptane and 2.75 mL of “fix mix” (4% paraformaldehyde in 0.1M Sorenson’s phosphate buffer: 0.133 M Na2HPO4, 0.133 M KH2PO4, pH 7.2). Fixed embryos were then transferred to a 1:1 heptane:methanol solution, shaken for 20 sec, rinsed five times in fix mix, then fixed for another 90 min in fix mix. Embryos were then stored in 1% paraformaldehyde in 0.1M Sorenson’s phosphate buffer overnight.

The following steps for cryo-sectioning and labeling were modified from a published protocol (*67*). After fixation embryos were washed twice with 0.15M glycine in PBS, then incubated in 1%, 5% and 12% gelatin in PBS at 37° C for 20 min each, and embedded in 12% gelatin. Embedded embryos were then cryo-protected in 2.3M sucrose in PBS overnight. Embryos were then frozen on aluminum pins by plunging into liquid nitrogen. Sections were cut at –100°C at a thickness of 75nm using a Leica Ultracut UCT ultramicrotome with a cryo-chamber attachment. Sections were picked up with a loop using a 1:1 mixture of 2% methylcellulose:2.3M sucrose. Sections were placed on Formvar coated nickel grids.

For immunolabeling, grids were floated on drops of PBS for 10 min to rehydrate sections, and then floated on drops of 0.15M glycine in PBS for 10 min to quench aldehydes. Sections were blocked with 5% cold-water-fish gelatin in PBS for 30 min, then incubated in primary antibody, guinea pig anti-SMG (1:40), rabbit anti-VAS diluted (1:20) in 1% cold-water-fish gelatin containing 0.1% Tween in PBS for 1 hr. Sections were then washed five times for 2 min each in PBS, followed by incubation with secondary antibody gold conjugate (goat anti-guinea pig, 10 nm gold, goat anti-rabbit 15 nm gold) diluted 1 in 10 in 1% cold-water-fish gelatin, 0.1% Tween in PBS for 1 hr. Sections were then washed five times for 2 min each in PBS, and fixed for 10 min with 2% glutaraldehyde in phosphate buffer; then washed twice for 2 min each in PBS, followed by five times 2 min each in dH2O. Sections were then incubated on drops of 2% methylcellulose, 0.4% uranyl acetate for 10 min. Grids were picked up in loops and the excess methylcellulose was wicked off. Sections were examined using an FEI Tecnai 20 transmission electron microscope. Images were captured using a Gatan Dualview digital camera.

### Fluorescence recovery after photobleaching (FRAP)

Embryos were mounted on Greiner Lumox culture dishes (Sigma-Aldrich Cat# Z376744) as described above. Samples were photobleached using an argon laser attached to the WAVEFX spinning disk (Quorum Technologies). A small ROI approximately the size of the germ granule was selected for photo bleaching. Images were captured using the 40x/NA 1.4 oil immersion lens (Carl Zeiss, Inc.), an EM charge-coupled camera (Hamamatsu Photonics), and Volocity Imaging software (Perkin Elmer). Samples were photobleached for 2 sec and images were collected at two frames/sec for a total of 4 min. Images were collected in a single Z-plane before, during and after photobleaching.

### Three-dimensional reconstruction and counting of PGCs and PH3-labeled nuclei

Series of confocal sections through the PGCs at the posterior pole were collected using an Olympus IX81 inverted fluorescence microscope equipped with a Hamamatsu Back-Thinned EM-CCD camera (9100-13) and Quorum spinning disk confocal scan head. Wild-type controls were at Stage 4 to 5 (nuclear cycle 12-14): 11.5% were in nuclear cycle 12, 68% in nuclear cycle 13, and 20.5% in nuclear cycle 14. Since the nuclear cycles cannot be assessed in *smg* mutant embryos (*23*), they were analyzed 2-3 hr after egg deposition.

For counting the number of PGC’s in the *osk*^2xSRE(+)^ and *osk*^2xSRE(–)^ transgenic lines in *osk*^*0*^*/Df(osk)* background, and in the *bru1*^*5xSRE(+)*^ and *bru1*^*5xSRE(–)*^ transgenic lines, a Leica SP8 inverted scanning confocal microscope using a 63x/NA 1.4 immersion oil objective was used to aquire confocal sections through the PGCs at the posterior pole of embryos in late NC 14. The LAS X (Leica Application Suite X) was used to examine images slice by slice and the annotation tool was used to mark and count PGCs.

### Western blotting

Dechorionated embryos, as described above, were counted and homogenized in lysis buffer: 150 mM NaCl, 50 mM Tris pH 7.5, 1 mM DTT, 1x protease inhibitor EDTA-Free cocktail (Roche). For the detection of transgenic OSK protein, a modified protease inhibitor cocktail (*68*) was added to the lysis buffer at the following concentrations: 0.2 mM benzamidine, 2 mM PMSF, 10 μM leupeptin, 10 μM pepstatin A, 4 μM chymostatin, 1 μM aprotinin. Extracts were boiled for 5 minutes in 2x SDS-PAGE loading buffer, and separated by 8% SDS-PAGE. Proteins were transferred to PVDF membranes (Immun-Blot PVDF, Bio-Rad, 1620177). Blots were blocked in 2% skim milk in PBST (PBS + 0.1% Tween 20) for 1 hr, and incubated at 4°C overnight with primary antibodies diluted in 1% BSA in PBST. Primary antibodies used were rabbit anti-BRU1 polyclonal (1:5000; a gift of Paul Macdonald, Austin, Texas), mouse anti-HA (1:10,000; Sigma clone HA7, H9658), mouse anti-Tubulin (1:3,000; Sigma clone B-5-1-2, T5168), and mouse anti-b-Actin (1:1,000; Abcam, AB8224).

After incubation with primary antibody, blots were washed 3 times for 10 min each with PBST at room temperature with rocking, then incubated at room temperature for 1 hr with HRP-conjugated secondary antibody diluted in 1% BSA in PBST (1:5000; either of Peroxidase-Affinipure goat anti-mouse IgG (H+L) or goat anti-rabbit IgG (H+L); Jackson Immunoresearch, Cat#115-035-003 and 111-035-114, respectively). Blots were washed again 3 times for 10 min each and developed using Immobilon Luminata Crescendo Western HRP substrate (Millipore, Cat#WBLUR0500). Western blots were imaged and quantified using ImageLab (Bio-Rad).

### Single-molecule fluorescence *in situ* hybridization (smFISH)

smFISH was performed as previously described (*30*). Probe sets with 2 nt spacing complementary to *nos* (CG5637; 63 oligos), *Hsp83* (CG1242; 48 oligos) or *bru1* (CG31762; 48 oligos) were designed using the Stellaris Probe Designer. Custom 20 nt oligonucleotides with a 3’NHS ester modification were obtained from Biosearch Technologies, conjugated to either Atto 565 (Sigma) or Atto 633 (Sigma) dye and purified via HPLC as described (*69*). Embryos were mounted in Prolong Diamond media (ThermoFisher P36965) and cured for 3- to-4 days at room temperature prior to imaging.

For temporal analysis of *osk* mRNA (Figure S2), imaging was performed using a Leica SP5 laser scanning microscope with a 63x/1.4 NA oil immersion objective and GaAsP “HyD” detectors in standard detection mode. All imaging parameters were kept identical within each experiment. For quantification of total intensity, z-series with a 2 µm step size were used to capture the germ plasm-localized signal. Embryos were staged using DAPI to determine nuclear cycle.

For colocalization of RNAs with Venus-SMG (Figure S4), imaging was performed using a 60x/1.4 NA oil immersion objective on a Nikon A1-RS laser-scanning confocal microscope, at 2x optical zoom, with pixels of 102 × 102 nm. In each experiment, laser power and gain were adjusted to avoid signal saturation while maximizing separation of signal and noise. Confocal sections were acquired using 16x line averaging. Embryos selected for imaging were oriented such that the germ plasm was facing the cover slip. To minimize fluorescent signal distortion, all images were taken within 5 µm of the embryo cortex. Venus-SMG was detected by direct GFP fluorescence.

### Quantification and Statistical Analysis

#### Analysis of SMG particle movement

SMG particle movement was measured using the MetaMorph software package (Universal Imaging, Downington, Pennsylvania). Individual particles that remained visible for four consecutive frames were tracked. The particle velocity was calculated by measuring the distance traveled by the particle divided by the transit time. The directionality of each particle was noted as posterior if the particle moved towards the posterior pole. Among the 47 particles that were tracked, 28 particles were directed towards the posterior while 19 particles moved towards the anterior of the embryo.

#### Fluorescence recovery after photobleaching (FRAP)

Signal recovery was measured using ImageJ (National Institutes of Health) for germ granules that stayed in the focal plane for the duration of the image acquisition. These values were corrected for background by subtracting the mean fluorescence of a similar size area outside of the embryo (*Pi*). To correct for general bleaching of the embryo from imaging, fluorescence was measured for a large area covering the entire pole cell or regions of apical cytoplasm, corrected for the background (*Bi*) as above. The corrected fluorescence intensity was measured by calculating *Pi* / *Bi* and normalized to the time point just before the bleaching, then plotted using Excel (Microsoft). Recovery rates were calculated from the slopes of the best-fit lines for the first 30 sec after photo-bleaching.

### Three-dimensional reconstruction and counting of PGCs and PH3-labeled nuclei

To count the total number of PGCs and the number with PH3-positive nuclei, the confocal stacks were imported into Volocity 5.5.0 software (PerkinElmer), for three-dimensional reconstruction. Embryos were then viewed in the ‘3D Opacity’ mode allowing one to freely turn, rotate and spin individual embryos in each plane in order to facilitate counting of PGCs, PH3-positive PGC nuclei, and to distinguish which PH3-positive signal was in the PGC nuclei versus in somatic nuclei (in *smg* mutants, where somatic nuclei continue to divide at these stages).

To count the number of PGC’s in the *osk*^2xSRE(+)^ and *osk*^2xSRE(–)^ transgenic lines in *osk*^*0*^*/Df(osk)* background, and in the *bru1*^*5xSRE(+)*^ and *bru1*^*5xSRE(–)*^ transgenic lines, the LAS X (Leica Application Suite X) was used to look at images slice by slice and the annotation tool was used to count PGCs.

PGC counts were input into Excel and graphs were produced using GraphPad software (Prism) for Figures 3A, 5C and 7C. For Figure 3A, the Kruskal Wallis One-Way ANOVA followed by Dunn’s test for multiple comparisons, was used to calculate *P* values. For Figures 5C and 7C, the one-sided Wilcoxon rank sum test was used to calculate *P* values. *P* values are presented in the figures and figure captions.

### Quantification of *osk* mRNA in wild type and *smg*-mutant embryos

Image processing and analysis were done in FIJI. Z-projections were made using the “sum slices” function and the threshold was adjusted so that the entire localized signal was included. The total fluorescence intensity of the localized signal (integrated density function in FIJI) was then measured and used to quantify average total *osk* fluorescence intensity for each genotype and stage. Significance was assessed using one-way ANOVA followed by the Tukey posthoc test for comparison to NC≤9 within each genotype. *P* values are presented in the figure and figure caption.

### Co-localization quantification

In MATLAB, a region encompassing the germ plasm was specified with a manually drawn polygon. A spot detection algorithm then defined the X,Y coordinates of every focus of protein or mRNA in each channel, using intensity thresholds set manually to eliminate false positives. Distances were calculated from each point in one channel to the nearest point in the other channel and *vice versa*. These values were concatenated from a pool of embryos, binned, and displayed as histograms. Functional colocalization is defined as distances less than 300 nm, on the basis of average germ granule diameter (*70)*.

### Western blot imaging and quantification

Western blots were imaged and quantified using ImageLab (Bio-Rad). In Figures 5 and 7 the OSK and BRU1 protein bands, respectively, were normalized to the b-Actin loading control band, and the signal for 0-1 hr SRE(+) embryos for the first replicate was set to 1 (n = 5 for Figure 5C, n = 4 for Figure 7C). Means of the replicates are presented in these figures with error bars representing standard deviation (Wilcoxon rank sum test, one-tailed, unpaired). *P* values are presented in the figures and figure legends.

## Acknowledgments

We thank Paul Macdonald for providing *osk*^0^ and *Df(3R)osk* fly lines, anti-OSK and anti-BRU1 antibodies, as well as cloned *osk* genomic DNA and the *bru1* cDNA; Paul Lasko for providing the VAS-GFP transgenic flies, anti-VAS and anti-OSK antibodies; Andrew Wilde for advice on spinning disk microscopy; and Tony Harris for assistance with the FRAP experiments. Special thanks to the Hospital for Sick Children Imaging Facility and Thomas Hurd for assistance with confocal microscopy. Electron microscopy was conducted at the Hospital for Sick Children/Mt. Sinai Hospital Electron Microscopy Facility with the assistance of Robert Temkin. Extensive use was made during this study of the following resources: FlyBase and the Bloomington Drosophila Stock Center.

## Funding

Canadian Institutes for Health Research grant MOP-14409 (HDL) Canadian Institutes for Health Research grant PJT-159702 (HDL) National Institutes of Health grant R01 GM061107 (ERG) National Institutes of Health grant R01 GM067758 (ERG) National Institutes of Health grant R35 GM067758 (ERG) Hospital for Sick Children Research Training Centre postdoctoral fellowship (ALG) National Institutes of Health grant T32 GM007388 (WVIE)

## Author contributions

Conceptualization: NUS, ALG, HDL

Methodology: HDL, ALG, NUS, AK, CAS, ERG, WVIE

Investigation: ALG, NUS, AK, CAS, ERG, WVIE

Visualization: ALG, NUS, AK, ERG, WVIE

Supervision: HDL, ERG

Writing—original draft: HDL

Writing—review & editing: HDL, ERG, CAS

## Competing interests

All authors declare they have no competing interests.

## Data and materials availability

All data are available in the main text or the supplementary materials. All materials (Drosophila strains, plasmids, etc.) are available from the corresponding author upon request.

## Notes

### Competing Interest Statement

The authors have declared no competing interest.

