## Supplementary data for "Smaug regulates germ plasm synthesis and primordial germ cell number in Drosophila embryos by repressing the *oskar* and *bruno 1* mRNAs"

**This PDF file includes:**

Figs. S1 to S4
Table S1
Movies S1 to S3

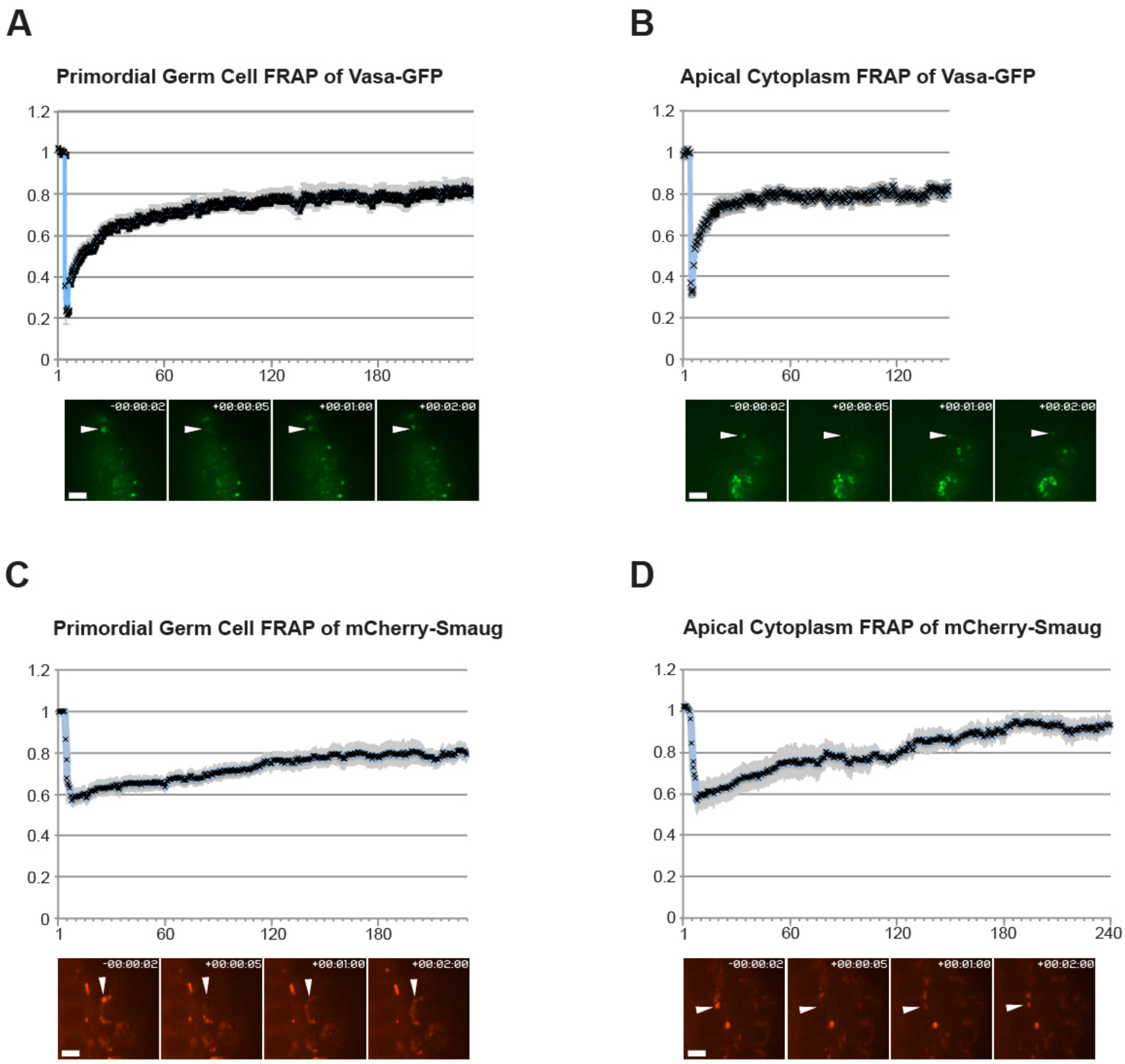


**Fig. S1, related to Fig. 2.**

**FRAP analysis of SMG and VAS protein in the germ granules.** (A) FRAP of VAS-GFP in germ granules in PGCs. (B) FRAP of VAS-GFP in germ granules that are left behind in soma. (C) FRAP of mCherry-SMG in germ granules in PGCs. (D) FRAP of mCherry-SMG in germ granules that are left behind in soma. The mobile fractions were 55% (SMG in PGC granules), 70% (VAS in PGC and left-behind granules), and 90% (SMG in left-behind granules). The mCherry-SMG transgene insertion was on the second chromosome (see Key Resources Table): *P{*$w^+$*[mCherry-SMG]:7}*.

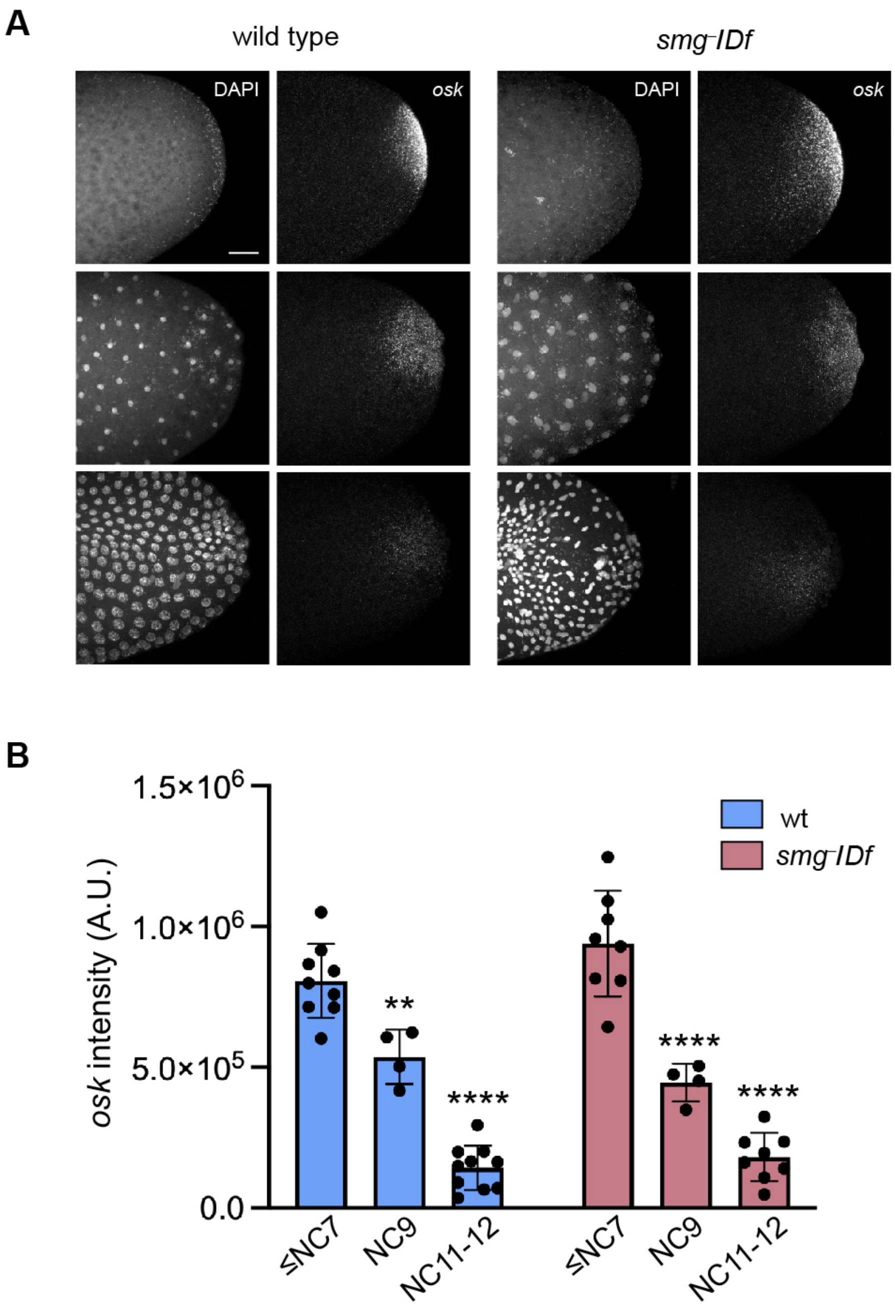


**Fig. S2, related to Fig. 4.**

***osk* mRNA levels in the germ plasm are not altered in *smg* mutants. (A)** Confocal z-series projections of wild-type or *smg*-mutant (genotype: $smg^1$/*Df(3L)Scf-R6*) embryos prior to pole cell budding (top panels), during pole cell budding (middle panels), and after pole cell formation (bottom panels). *osk* mRNA was detected by smFISH and DNA was visualized using DAPI. Note that *smg* mutants show nuclear cycle defects after NC 10 as seen in the bottom righthand DAPI panel. Scale bar = 25 μm. **(B)** Quantification of the average total *osk* fluorescence intensity in the germ plasm in embryos at the indicated stages. Data points represent individual embryos, mean ± s.d. is shown; $**P < 0.01$, $****P < 0.0001$ for comparison to NC≤9 within each genotype, as determined by one-way ANOVA and Tukey posthoc test.

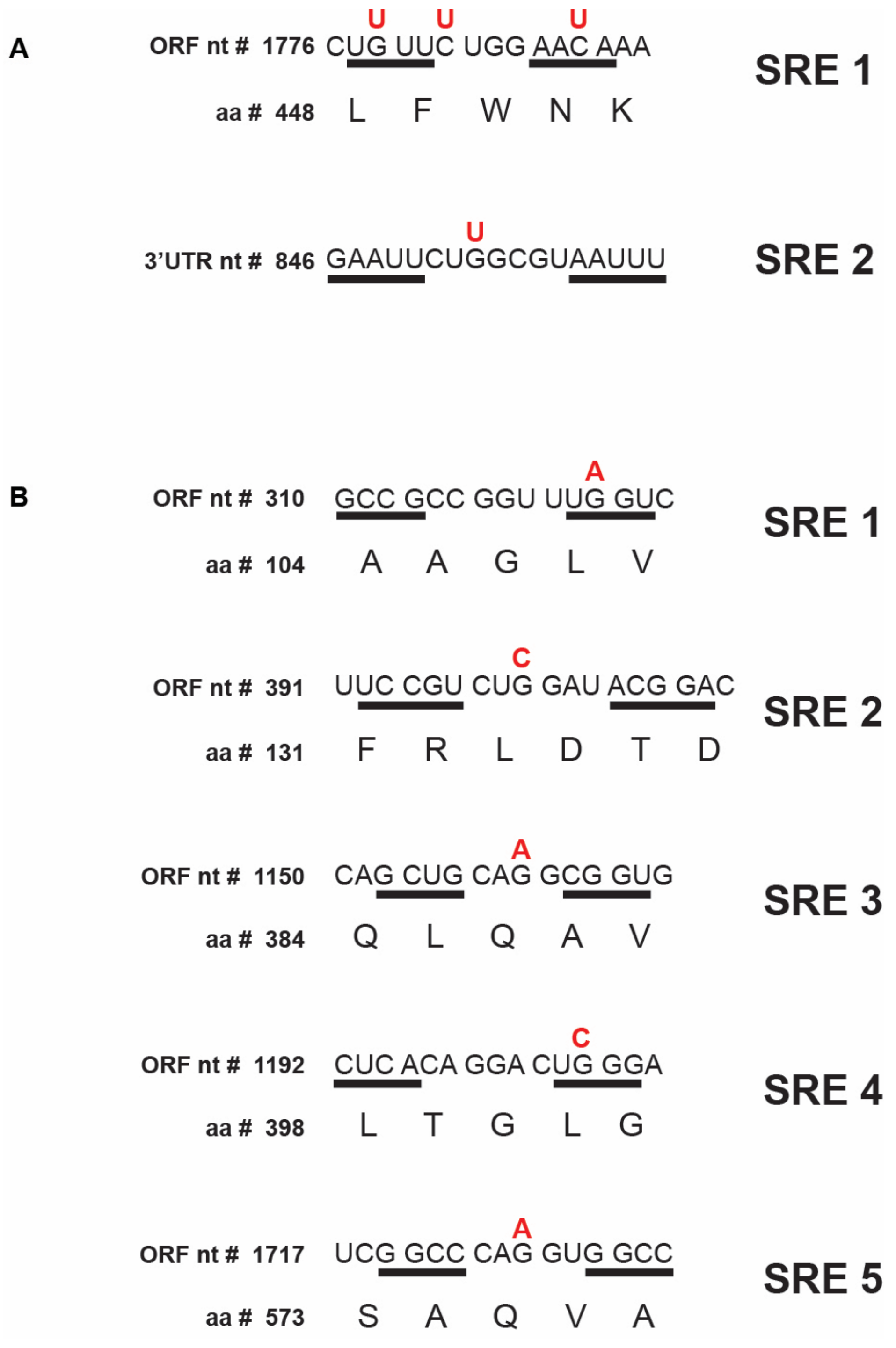


**Fig. S3, related to Figs. 5 and 7.**

**Wild-type and mutant versions of the SREs in the *osk* and *bru1* transgenic mRNAs. (A)** Sequence of the SREs in mRNAs encoded by the $osk^{2xSRE1(+)}$ and $osk^{2xSRE1(–)}$ transgenes. **(B)** Sequence of the SREs in the mRNAs encoded by the $bru1^{5xSRE(+)}$ and $bru1^{5xSRE(–)}$ transgenes. Nucleotides predicted to form the SRE stems are underlined. Mutated nucleotides are shown in red. In all cases where the SREs reside in the open reading frame, the encoded amino acids were unchanged by the mutations.

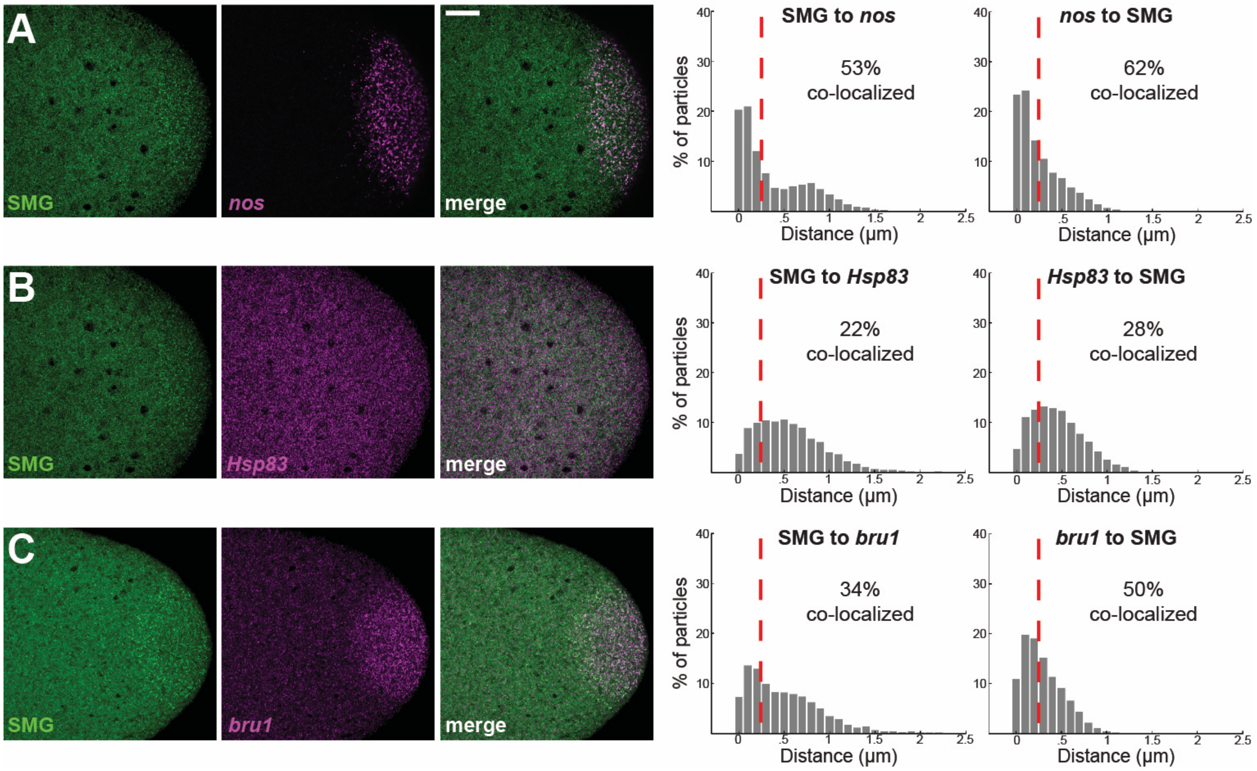


**Fig. S4, related to Fig. 6.**

**SMG colocalizes with *nos* and *bru1* but not *Hsp83* mRNA in mature germ plasm. (A-C)** Confocal sections of the posterior region of 0-to-2 hr old embryos expressing Venus-SMG (green) with anterior to the left and dorsal towards the top of the page. *nos* **(A)**, *Hsp83* **(B)** and *bru1* **(C)** mRNAs (magenta) were visualized with complementary smFISH probes. The *nos* probes (Atto 633-labeled) and *Hsp83* probes (Atto 565-labeled) were applied to the same Venus-SMG embryos. The same embryo is shown in **(A)** and **(B)**. Scale bar: 15 µm. Nearest-neighbor quantification of co-localization between SMG and *nos*, *Hsp83* or *bru1* transcripts is shown at the right (n = 11, 10 and 13 embryos, respectively). The dashed red line indicates the threshold of 300 nm used to define co-localization

| Genotype of female parent† | Number of embryos | PGC number | | Phosphohistone H3 positive nuclei | |
|---|---|---|---|---|---|
| | | Mean | S.D. | Mean | S.D. |
| **Wild-type** *$w^{1118}$* | 44 | 31.8 | 7.0 | 2.5 | 2.3 |
| ***smg* mutant** | | | | | |
| *$smg^{1}/Df$* | 38 | 38.0 | 8.4 | 1.0 | 1.2 |
| *$smg^{1}/smg^{47}$* | 42 | 37.5 | 9.2 | 0.7 | 1.3 |

† All genotypes carried the VAS-GFP transgene to mark the PGCs. *Df* is a deletion that removes the *smg* gene: *$smg^{1}/Df(3L)Scf\text{-}R6$*. S.D.: standard deviation.

**Table S1, related to Fig. 3.**

**PGC and PH3 numbers in wild type and in *smg* mutants**
